# Distinct hyperactive RAS/MAPK alleles converge on common GABAergic interneuron core programs

**DOI:** 10.1101/2022.08.04.502867

**Authors:** Sara J Knowles, April M Stafford, Tariq Zaman, Kartik Angara, Michael R Williams, Jason M Newbern, Daniel Vogt

**Affiliations:** School of Life Sciences, Arizona State University, Tempe, AZ, 85287, USA; Department of Pediatrics and Human Development, Michigan State University, Grand Rapids, MI, 49503, USA; Neuroscience Program, Michigan State University, East Lansing, MI, 48825, USA

## Abstract

RAS/MAPK gene dysfunction underlies various cancers and neurocognitive disorders. While the role of RAS/MAPK genes have been well studied in cancer, less is known about their function during neurodevelopment. There are many genes that work in concert to regulate RAS/MAPK signaling, suggesting that if common brain phenotypes could be discovered they could have a broad impact on the many other disorders caused by distinct RAS/MAPK genes. We assessed the cellular and molecular consequences of hyperactivating the RAS/MAPK pathway using two distinct genes in a cell type previously implicated in RAS/MAPK-mediated cognitive changes, cortical GABAergic interneurons. We uncovered some GABAergic core programs that are commonly altered in each of the mutants. Notably, hyperactive RAS/MAPK mutants bias developing cortical interneurons towards those that are somatostatin+. The increase in somatostatin+ interneurons could also be induced by elevated neural activity and we show the core RAS/MAPK signaling pathway is one mechanism by which this occurs. Overall, these findings present new insights into how different RAS/MAPK mutations can converge on GABAergic interneurons, which may be important for other RAS/MAPK genes/disorders.

## Introduction

Cellular signaling via the RAS/MAPK cascade is a critical regulator of multiple cellular and molecular developmental milestones (Seger and Krebs, 1995; Sun et al., 2015; Waltereit and Weller, 2003). These signaling events translate various extracellular cues to downstream effectors in both the cytosol and nucleus to impact cell proliferation, migration, morphology and synapse maturation/plasticity. Importantly, mutations in RAS/MAPK genes underlie a family of neurodevelopmental syndromes with an elevated risk of autism spectrum disorder (ASD) and cancer (Adviento et al., 2014; Hoshino et al., 1999; Vithayathil et al., 2018). Several animal studies have led to insights into how dysfunctional RAS/MAPK genes impact brain function, reviewed in (Gutmann et al., 2012; Hebron et al., 2022; Kang and Lee, 2019). However, a more in-depth investigation of specific brain cell types at the cellular and molecular level that may underlie the cognitive symptoms is needed. Common phenotypes between these disorders could have major implications for future therapeutics.

Earlier studies examining the RAS/MAPK pathway inhibitor, *Nf1*, suggested that GABAergic dysfunction could be a key factor in the cognitive changes associated with RAS/MAPK disorders (Costa et al., 2002; Cui et al., 2008). Recent studies identified specific cellular and molecular consequences of RAS/MAPK hyperactivation in GABAergic cortical interneurons (CINs), including the loss of parvalbumin (PV)+ CINs and a decrease in LHX6 (Angara et al., 2020; Holter et al., 2021; Omrani et al., 2015). LHX6 is a cardinal transcription factor that is necessary for the emergence of CIN populations from the medial ganglionic eminence (MGE) (Liodis et al., 2007; Vogt et al., 2014; Zhao et al., 2008). MGE-derived CINs primarily express either PV or somatostatin (SST) (Liodis et al., 2007; Zhao et al., 2008), constitute ~70% of forebrain CINs and are necessary players in brain microcircuit function and disease (Marín, 2012; Wonders and Anderson, 2006). A critical gap in knowledge is how distinct GABAergic CINs become fated to attain their unique molecular, morphological and electrophysiological signatures (Hu et al., 2017a; Lim et al., 2018; Mayer et al., 2018; Wamsley and Fishell, 2017). Whether cellular events, particularly RAS/MAPK signaling could be involved, has not been thoroughly explored. This is an important developmental question, as the PV and SST interneuron types are derived from the same progenitor cells in the embryonic medial ganglionic eminence (MGE) (Hu et al., 2017a; Wamsley and Fishell, 2017; Wonders and Anderson, 2006), yet mature into distinct cell types in mice. One hypothesis of how distinct properties arise is through engagement of activity dependent processes as CINs integrate into their respective target locations (Close et al., 2012; De Marco García et al., 2011; Denaxa et al., 2012; Wamsley and Fishell, 2017). Since RAS/MAPK signaling is elevated by neural activity (Adams and Sweatt, 2002; Thomas and Huganir, 2004; West et al., 2001), it is possible that activity-dependent recruitment of RAS/MAPK impacts the development of GABAergic interneurons via changes in core transcriptional programs necessary for their development. Despite these observations, no one has tested that these observations converge in CINs.

We thus investigated whether core GABAergic and CIN developmental programs were altered in two distinct genetic animal models that lead to hyperactive RAS/MAPK signaling, building upon recent work that examined how hypofunction of the RAS/MAPK pathway impacts development (Knowles et al., 2022). While mutations in RAS/MAPK signaling genes are implicated in cognitive changes in the RASopathies, there is substantial variability between individuals, potentially due to their specific gene mutation and/or hierarchy of the gene product in the signaling pathway (Adviento et al., 2014). Despite these challenges, common phenotypic changes shared between different RAS/MAPK mutants may also exist and could be a fundamental inroad to treat overlapping symptoms in RASopathies. To uncover these features, we assessed *Nf1* loss of function and *bRaf* constitutively active (ca) genetic mouse models in CINs, with the goal of identifying what common changes occur when RAS/MAPK signaling was amplified.

We uncovered RAS/MAPK-alterations in CINs impacting core developmental genes involved in cell fate and function. Hyperactive RAS/MAPK gene mutants resulted in a bias towards somatostatin (SST) expressing cells with correlative physiological properties at the expense of parvalbumin (PV) expressing CINs. We also found that neuronal activity-induced RAS/MAPK signaling is one way in which SST expressing CINs are selectively biased, potentially bridging several known observations about neural activity and its role in recruiting RAS/MAPK signaling (Tyssowski et al., 2018; Wiegert and Bading, 2011) as well as growth factor and activity-induced SST expression (Tolon et al., 1994; Zeytin et al., 1988). These results suggest that a common GABAergic phenotypic program is altered in hyperactive RASopathies and that RAS/MAPK signaling is one conduit for how extracellular cues/cellular signaling can influence MGE molecular properties.

## Results

### *Nf1* and *bRaf^ca^* mutants exhibit similar decreases in PV but distinct changes to SST CINs by adult ages

We used a genetic approach to manipulate different RAS/MAPK genes, first comparing *Nf1* loss with *bRaf* constitutively active (ca) mutants; each results in hyperactivation of the MAPK signaling cascade. The function and stratification of these and other RAS/MAPK proteins are shown in Supp. Fig 1. This approach allowed us to discern phenotypes resulting from *Nf1* deletion (upstream inhibitor of pathway), which regulates multiple signaling cascades, versus selective hyperactivation of the RAS/MAPK pathway, via downstream *bRaf* activation. Cre-dependent *bRaf^ca^* (Urosevic et al., 2011) or *Nf1* floxed mice (Zhu et al., 2001) were crossed with *Nkx2.1-Cre* (Xu et al., 2008) and *Ai14* alleles (Madisen et al., 2010) to generate WT, *Nf1* conditional knockouts (cKO) and hemizygous *bRaf^ca^* embryos that express tdTomato in Cre-recombined cells.

We first needed a way to compare these two gene manipulations in CINs of young adult mice. However, two issue had to be managed. *Nkx2.1-Cre* induced recombination resulted in no live *Braf^ca^* pups, precluding adult assessments and *Nf1* mutants exhibit elevated numbers of premature oligodendrocytes (Angara et al., 2020). To navigate these obstacles, we used an MGE cell transplantation approach that has been used to assess molecular and cellular phenotypes of mature CINs in vivo from mutant mice that exhibit premature lethality (Vogt et al., 2014), thus solving the *bRaf^ca^* lethality. In addition, CINs are unique in their ability to disperse and migrate once transplanted into the brain (Alvarez-Dolado et al., 2006), allowing us to physically separate CINs from oligodendrocytes in vivo. To this end, embryonic day (E)13.5 MGE cells were collected from *Nkx2.1-Cre; Ai14* embryos that were WT, *Nf1* cKO or *Braf^ca^*, transplanted into postnatal day (P)2 WT neocortices and allowed to develop in vivo for 35 days (Schema, Figure 1A).

**Figure 1:**
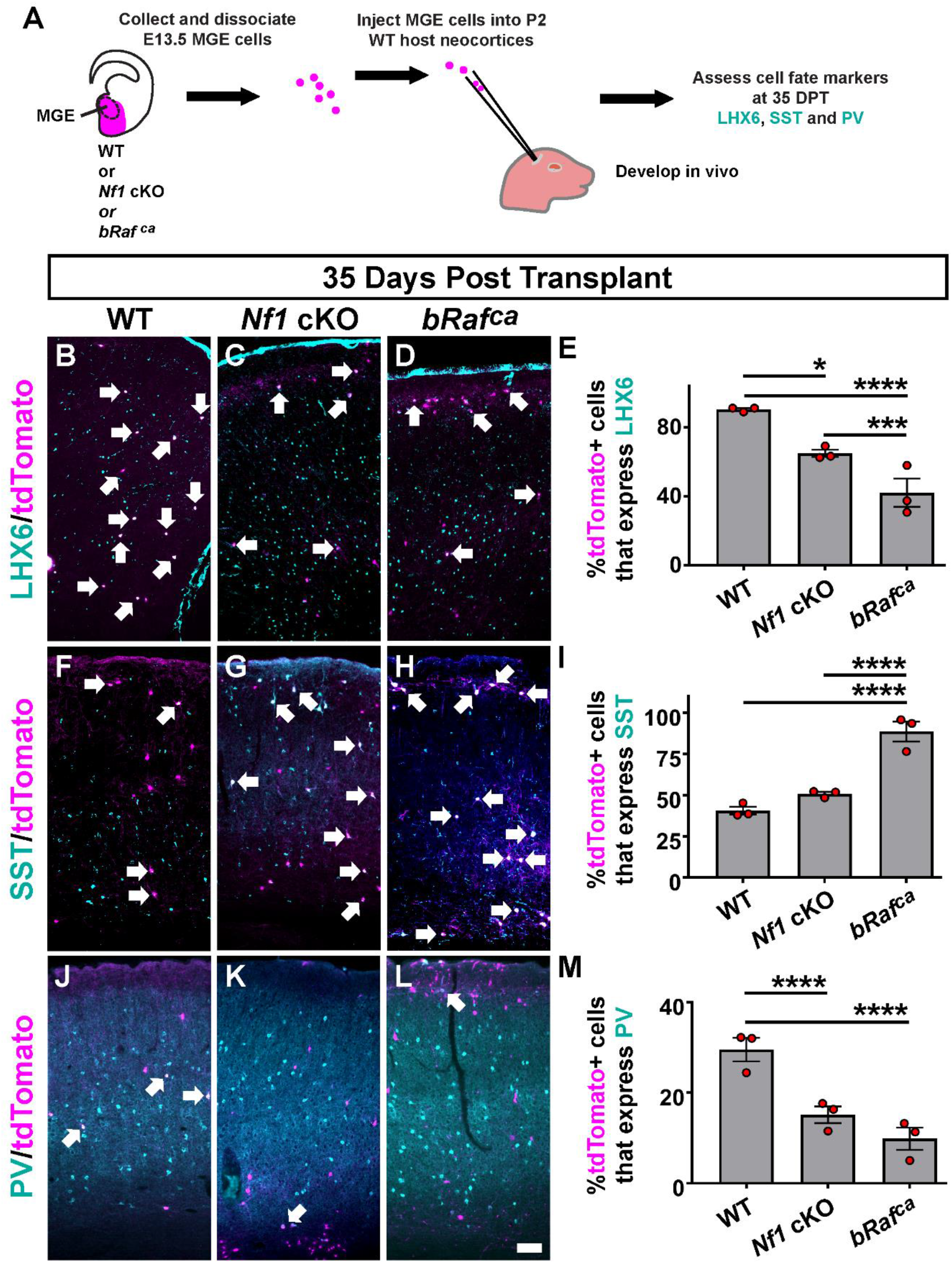
*Nf1* and *Braf* MGE transplants reveal altered LHX6, SST and PV expression by mature ages: (A) Schema depicting the MGE cell transplantation assay. Briefly, E13.5 MGE progenitors were harvested, dissociated and then injected into the neocortex of a WT host neonatal mouse. The cells developed and matured in vivo and were then assessed for molecular markers 35 days post-transplant (DPT). Transplanted and mature WT, *Nf1* cKO or *bRaf^ca^* CINs were then assessed for the proportion of MGE transplanted CINs that express LHX6 (B-E), SST (F-I) or PV (J-M), revealing decreased LHX6 and PV expression in both mutant CINs and a unique increase in SST expression in the *bRaf^ca^* CINs. Data are expressed as the mean ± SEM, n = 3 for each group. * p<0.05, *** p<0.001 and **** p<0.0001. Scale bar in (L) = 100μm.

The transplanted cells expressed tdTomato and were co-labeled for LHX6, SST or PV (Figure 1 B-D, F-H, J-L), allowing us to assess the proportion of MGE-lineage transplanted cells that expressed each marker after their development and maturation in vivo. The percentage of *Nf1* cKO and *Braf^ca^* tdTomato+ cells that expressed LHX6 was decreased by 28% and 50%, respectively, compared to WTs, providing support that this molecular phenotype is cell autonomous and shared between the mutants (Figure 1E; WT vs. *Nf1* cKO p = 0.04, WT vs. *Braf^ca^* p < 0.0001, *Nf1* cKO vs. *Braf^ca^* p = 0.0002). Of note, we did detect tdTomato+ oligodendrocytes in *Nf1* cKO transplants but they remained at the injection site; tdTomato+ oligodendrocytes were not found in *bRaf^ca^* transplants (data not shown).

We next examined the expression of SST in the transplanted cells. In agreement with our previous studies, the proportion of *Nf1* cKO cells at this mature age that expressed SST was similar to WTs (Figure 1I) (Angara et al., 2020; Holter et al., 2021). In contrast, most of the *Braf^ca^* cells expressed SST at high levels (Figure 1I; WT and *Nf1* cKO vs. *Braf^ca^* p < 0.0001). Finally, we determined the proportion of transplanted cells that expressed PV. Both the *Nf1* cKOs and *Braf^ca^* mutants had decreased expression of PV, 48% and 70%, respectively (Figure 1M; WT vs. *Nf1* cKO and *Braf^ca^* p < 0.0001). Overall, each mutant exhibited alterations in CIN markers with the more pronounced phenotypes observed in *bRaf^ca^* mutants.

### Postmitotic depletion of *Nf1* leads to a reduction in LHX6

We next tested if the loss of LHX6 was due to alteration in MGE progenitor cells or if this was a postmitotic phenomenon. To this end, we crossed both *Nf1^Flox^* and *bRaf^ca^* mice to *Lhx6-Cre* mice, to deplete the genes at a later developmental stage, as cells are becoming postmitotic. Unfortunately, we were not able to collect live *Nf1* cKO or *bRaf^ca^* progeny at postnatal stages, likely due to *Lhx6-Cre* recombination in blood vessels (Fogarty et al., 2007). However, we acquired viable *Nf1* cHet mice, which survived to P30, to assess LHX6 protein expression. We found a ~47% reduction of LHX6 expression in *Lhx6-Cre; Nf1* cHets compared to WTs (Supp. Fig. 2A-C, p = 0.004). These data indicate that reduced Nf1 in postmitotic neurons can suppress LHX6 expression and this phenotype is not due to disruption of progenitor MGE cell biology.

### *bRaf^ca^* mutant CINs exhibit a reduction in action potential spiking kinetics

The elevated ratio of SST+ to PV+ CINs in *bRaf^ca^* mutants (Figure 1) suggested that these mutants may exhibit a shift in CIN properties towards a SST-like CIN at the expense of the PV group. SST+ and PV+ CINs have distinct electrophysiological properties. SST+ CINs are mostly regular spiking and exhibit spike amplitude adaptation over time, while putative PV+ CINs are fast spiking with little to no adaptation (Halabisky et al., 2006; Hu et al., 2014; Kepecs and Fishell, 2014). Thus, if hyperactive *bRaf* resulted in a more general shift in cell properties, we hypothesized that mutant CINs would also exhibit a loss of faster spiking properties. Current clamp recordings were performed in S1 region of the neocortex to measure spontaneous and evoked activity; example transplanted cells are shown (Figure 2A, A’, A”).

**Figure 2:**
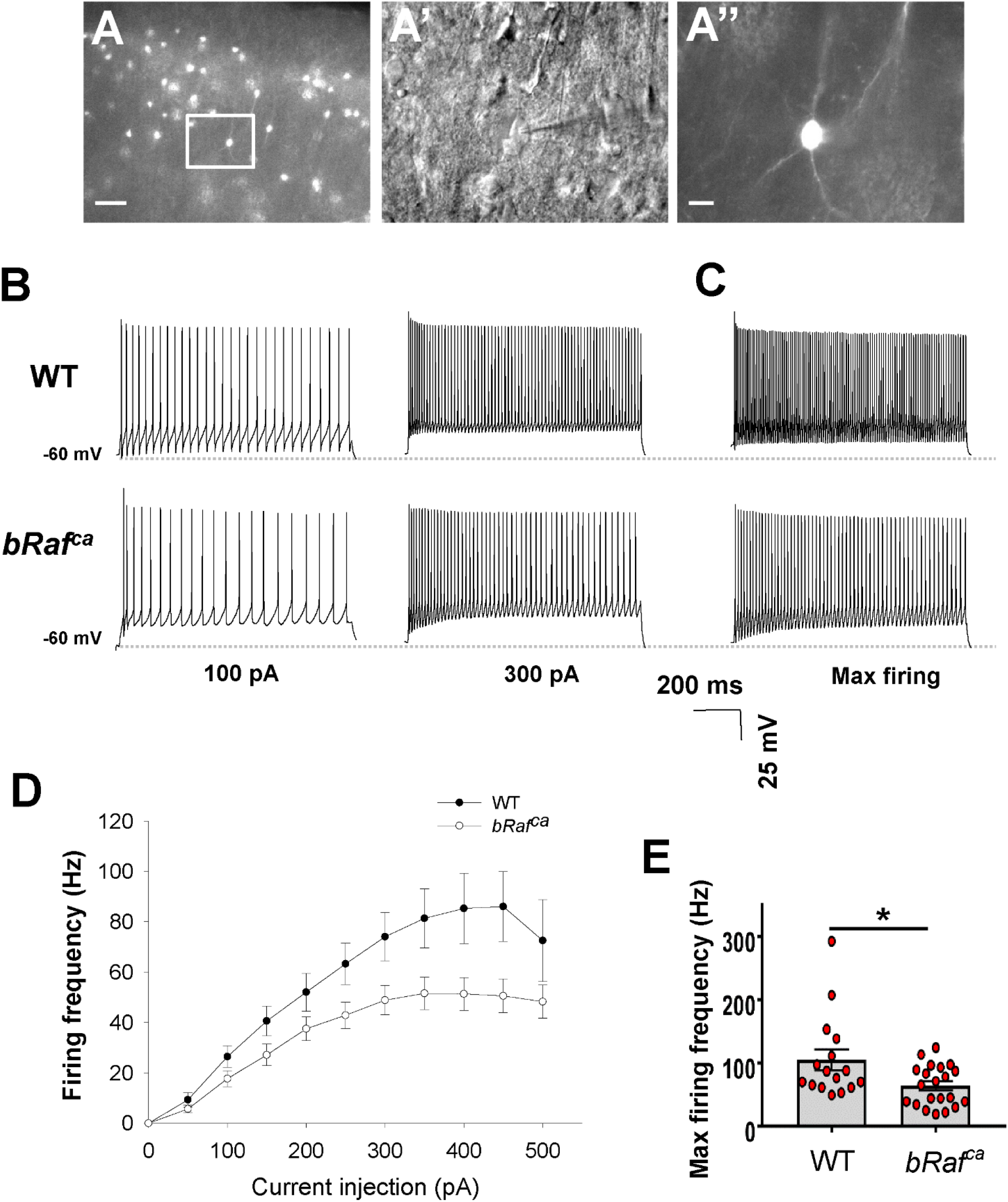
*bRaf^ca^* mutant CINs exhibit reduced action potential firing frequency: (A, A’) Images showing S1 region of the cortex (S1) with tdTomato+ transplanted CINs. (A’’) differential interference contrast (DIC) images of a patched CIN. (B) Representative traces showing neuronal firing in response to 100 and 300 pA current injections in WT (upper traces) and *bRaf^ca^* (lower traces). (C) Representative traces during maximal firing. (D) Quantification of evoked firing frequency in WT and *bRaf^ca^* CINs at different current injections, Two-way repeated ANOVA for firing properties, comparison at different level: Groups (F (1,35) = 6.702; p 0.014), current increment (F (1,35) = 31.555; p 0.001), and interaction of current and groups: (F (1,35) = 1.847; p = 0.05). (E) Quantitative analysis of maximal firing frequency between WT and *bRaf^ca^* CINs. Data are presented as the mean ± SEMs. * p<0.05. The horizontal dotted line in the traces indicates −60 mV. Abbreviations: milliVolts (mV) and picoAmps (pA). Scale bars in (A) = 50μm and (A’’) = 10μm.

We assessed whether action potential spiking was different between WT and *bRaf^ca^* cells. Example traces of spiking are shown for 100 pA and 300 pA current injections between genotypes (Figure 2B) as well as during maximum firing (Figure 2C). Consistent with our hypothesis, a two-way repeated-measures ANOVA revealed a significant effect of groups (Figure 2D) (F (1,35) = 6.702; p 0.01), current increment (F (1,35) = 31.555; p 0.001), and trending interaction of current and groups: (F (1,35) = 1.847; p = 0.05); action potential amplitude for both groups was similar. Finally, maximum evoked spike frequency was significantly reduced in *bRaf^ca^* CINs (Figure 2E, p = 0.02). These data support that *bRaf^ca^* mutants can promote CIN electrophysiological properties towards lower action potential spiking frequencies.

We also assessed other passive and active properties of the transplanted CINs (Supp. Table 1). Many properties were not significantly changed, including membrane capacitance as well as resting and active membrane resistance. Importantly, mutant CINs mostly resembled WT cells, suggesting proper maturation. However, resting membrane potential was elevated in the *bRaf^ca^* transplanted cells (p = 0.006). Consistent with the decreased maximum firing frequency, we also noticed altered interspike interval (ISI) length in the mutant cells; the initial ISI in the mutants trended towards longer duration (n.s., p = 0.08) and the last ISI was significantly longer in the mutants (p = 0.04). We also examined the spike frequency adaption (ratio of last ISI to first) but this was not significantly different, probably due to long first ISIs in the mutants. Overall, *bRaf^ca^* mutant CINs have shifted dynamics that are more aligned with SST+/regular spiking/adapting CINs but may not exhibit a full shift in properties towards this group.

### Elevated SST expression is a common hyperactive RAS/MAPK phenotype

To assess whether elevated SST levels and/or numbers of cells are a common phenotype in hyperactive RAS/MAPK mutants, we first assayed SST protein in MGE primary neuronal cultures from E13.5 brains, aged 8 days in vitro (schema Fig. 3A). Both *Nf1* cKO and *bRaf^ca^* cultures exhibited an elevated percentage of SST+ CINs (Figure 3B-H p < 0.0001). Qualitative increases in total SST that filled *bRaf^ca^* mutant cells were also noted (Figure 3D), suggesting that SST protein expression is a shared feature of elevated RAS/MAPK activity.

**Figure 3:**
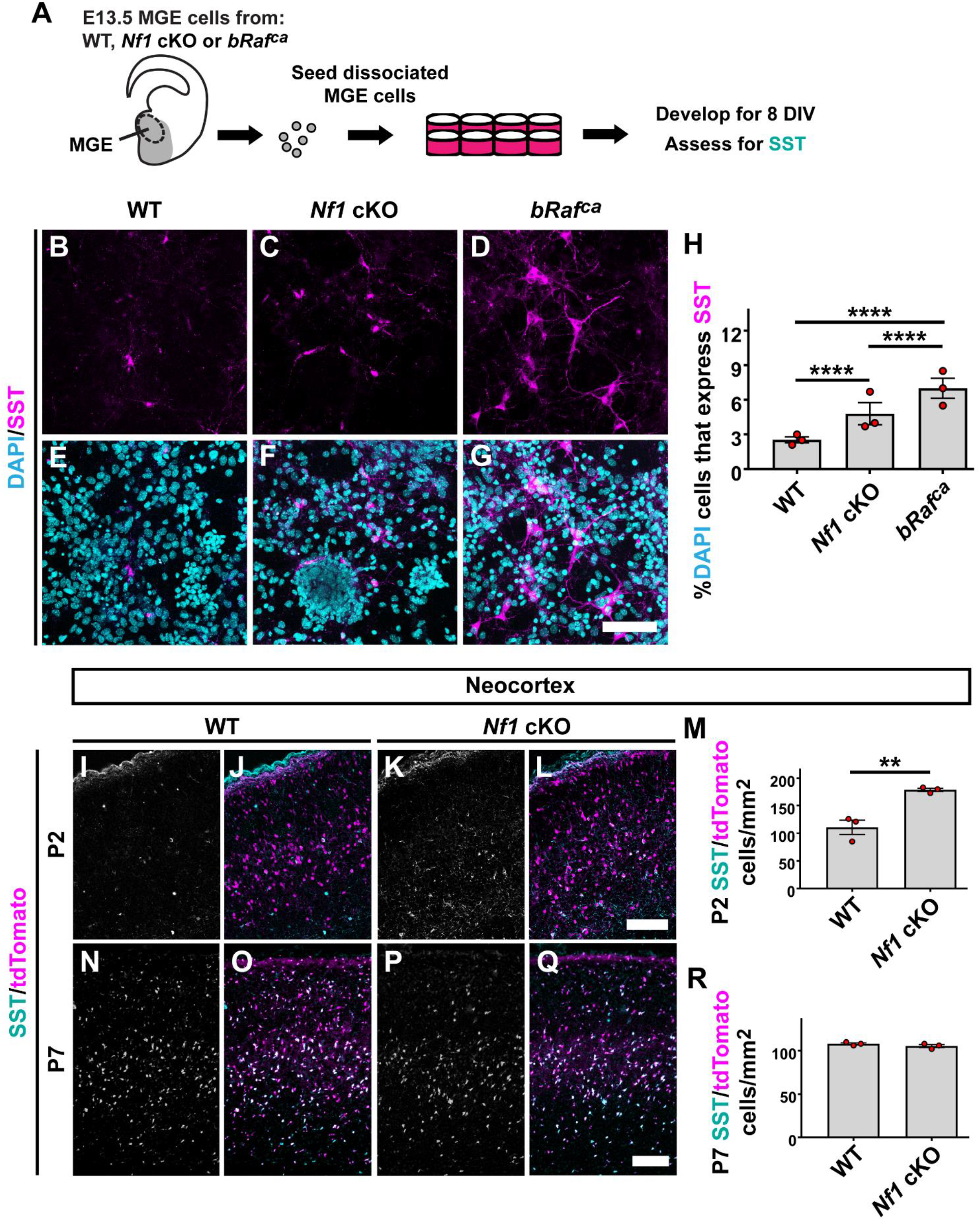
Elevated SST CINs are a common phenotype of *Nf1* and *bRaf* mutants in early development: (A) Schema depicting MGE primary culture procedures (gray area depicts *Nkx2.1-Cre* domain); E13.5 MGE progenitors dissociated and grown 8 days in vitro (DIV) before assessing SST levels. (B-G) images of SST and DAPI stained primary cultures after 8DIV. (H) Quantification of the proportion of DAPI+ cells that expressed SST; elevated SST numbers are found in both mutants. (I-L and N-Q) Neocortical images of CINs (tdTomato+) co-labeled for SST. (M and R) Quantification of tdTomato+/SST+ cells at P2 and P7, respectively, showing elevated levels early that normalize by P7. Data are expressed as the mean ± SEM, n = 3 for each group. ** p<0.01 and **** p<0.0001. Scale bars in (G) = 50μm and (L and Q) = 100μm.

We also examined SST expression at early postnatal stages to determine if the *Nf1* cKOs exhibited elevated SST expression in vivo. The previous primary culture experiments were aged in vitro to an equivalent age of postnatal day (P)2, thus we assessed SST levels at P2 in the neocortex and found ~62% increase in *Nf1* cKO CINs expressing SST (Figure 3I-3M p = 0.007), consistent with the primary cultures. By P7 there was no difference in SST expression between WTs and *Nf1* cKOs (Figure 3N-3R). Since both the *Nf1* cKO and *bRaf^ca^* embryos had elevated SST+ levels without changes in total tdTomato+ CINs (SJK and JMN, data not shown and (Angara et al., 2020)), we first concluded that hyperactive MAPK mutants have a developmental preference to bias MGE towards SST+ CINs.

The early developmental preference in the mutants to bias SST+ over PV+ CINs could explain the deficit in PV+ CINs at more mature ages. However, there are some discrepancies between different mutations; *bRaf^ca^* mutant CINs had elevated SST+ numbers but *Nf1* cKOs had normal levels at adult stages. The developmental stage between P2 and P7 for CINs is marked by programmed apoptosis (Southwell et al., 2012) which relies, in part, on RAS/MAPK signaling. To assess the influence of hyperactive MAPK mutations on cell death we performed MGE transplants of WT, *Nf1* cKO and *bRaf^ca^* MGE cells that developed for 13 days post-transplant and were then assessed during their peak apoptosis window (Southwell et al., 2012). We found less apoptosis in *bRaf^ca^* mutants (Supp. Fig. 3 *bRaf^ca^* vs. WT p = 0.006 and *Nf1* cKO p = 0.045). Thus, while *Nf1* cKO and *bRaf^ca^* CINs each exhibit increased SST expression early, the *bRaf^ca^* CINs partially elude programmed apoptosis during development, resulting the same loss of PV but differential SST ratios in mature cells.

### *Nf1* and *bRaf^ca^* mutations have unique and common effects on core MGE proteins

CIN development is regulated by well-defined transcription factors, though how these programs are influenced by activity and MAPK signaling is largely unknown. Thus, we asked if *Nf1* cKO and *bRaf^ca^* mutants altered core GABAergic and MGE-lineage programs in the embryonic forebrain. To this end, we focused on proteins involved in these programs in either *Nf1* cHet or cKOs as well as *bRaf^ca^* embryos. We chose E15.5 for assessment, as brains at this age have MGE-derived cells that are undergoing multiple developmental milestones, including continued propagation and migration throughout the cortex. Dissection of the brain (Schema, Figure 4A) was performed to remove hindbrain/midbrain structures while preserving forebrain.

**Figure 4:**
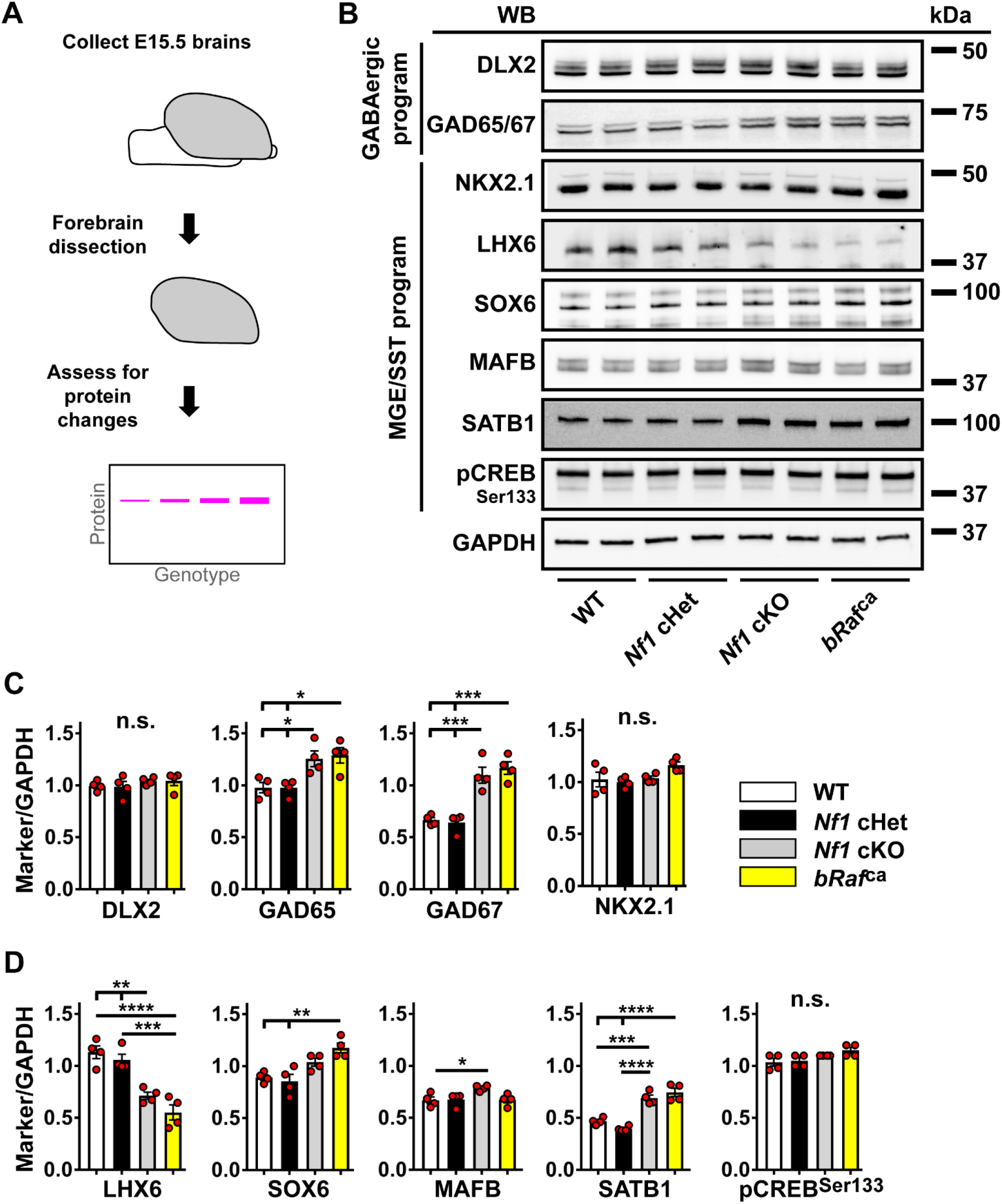
Biochemical assessment of GABAergic and MGE-lineage genes: (A) Schema to depict the collection of forebrain tissue for western blot analyses from E15.5 forebrains. (B) Western blots from representative pairs of genotypes that probed for GABAergic program genes and more specific MGE/SST program genes; SOX6 is the middle band in the image. GAPDH was used as a loading control. (C) Quantification of the band intensity divided by GAPDH band intensity for GABAergic and patterning markers. (D) Quantification of band intensities for more specific MGE/SST program markers. Data are expressed as the mean ± SEM, n = 4 for each group. * p<0.05, ** p<0.01, *** p<0.001 and **** p<0.0001. Abbreviations: (WB) western blot, (kDa) kiloDaltons.

Western blots for candidate proteins involved in the broad GABAergic program (DLX2 and GAD65/67) or MGE patterning (NKX2.1) were performed (Figure 4B). DLX2 and NKX2.1 levels were unchanged (Figure 4B and C). GAD65/67 levels were increased in *Nf1* cKO and *bRaf^ca^* brains (Figure 4B and C GAD65: WT and *Nf1* cHet vs. *Nf1* cKO p = 0.3, WT and *Nf1* cHet vs. *bRaf^ca^* p = 0.02; GAD67: WT vs. *Nf1* cKO p = 0.0006, WT vs. *bRaf^ca^* p = 0.0002, *Nf1* cHet vs. *Nf1* cKO p = 0.0004, *Nf1* cHet vs. *bRaf^ca^* p = 0.0001) suggesting a role for MAPK activity on the activity-dependent regulation of *Gad* genes (Hanno-Iijima et al., 2015). Uncropped images are shown in supplemental data.

### LHX6 is commonly downregulated in *Nf1* cKO and *bRaf^ca^* mutants

As expected, LHX6 protein was decreased in both *Nf1* cKOs and *bRaf^ca^* brains (Figure 4B and D WT vs. *Nf1* cKO p = 0.001, WT vs. *bRaf^ca^* p <0.0001, *Nf1* cHet vs. *Nf1* cKO p = 0.006, *Nf1* cHet vs. *bRaf^ca^* p = 0.0002); levels in *Nf1* cHets decrease at later ages (Supp. Figure 2 and (Angara et al., 2020)). Additionally, we assessed embryonic day E15.5 *bRaf^ca^* brains for LHX6 protein expression in *Nkx2.1-Cre*-lineage cells. While the cell density of *Nkx2.1-Cre*-lineage cells (tdTomato+) was not altered between genotypes (Supplemental Figure 4A, D, G, J, M), the proportion of tdTomato+ cells that co-labeled for LHX6 protein were only ~half as numerous in the *Braf^ca^* compared with littermate controls in the neocortex (Supplemental Figure 4B, C, E, F, H, I, K, L, N; p<0.0001). Thus, *bRaf^ca^* mutants exhibit an early loss of LHX6, more severe than *Nf1* cHet and cKO mutants.

### SATB1 is commonly upregulated in *Nf1* cKO and *bRaf^ca^* mutants

LHX6 can modulate the expression of several genes that may underlie increased SST in the mutants. To this end, we examined three markers known to be involved in the promotion of SST cell fate; SOX6, MAFB and SATB1 (Close et al., 2012; Denaxa et al., 2012; Hu et al., 2017b; Pai et al., 2019; Vogt et al., 2014). Mildly elevated MAFB was found in *Nf1* cKOs (Figure 4B, 4D p = 0.04), though not *bRaf^ca^* samples, suggesting a potential unique role for *Nf1* on the control of this MGE-lineage gene. SOX6 also had elevated expression within the *bRaf* but not *Nf1* mutants (Figure 4B, 4D WT vs. *bRaf^ca^* p = 0.005; *Nf1* cHet vs. *bRaf^ca^* p = 0.002). Surprisingly, pCREB was not altered in the mutants (Figure 4B, 4D), despite reported positive regulation by activity and RAS/MAPK signaling as well as its ability to directly transduce SST (Gonzalez and Montminy, 1989; Wu et al., 2001). However, the most striking change was the increase in SATB1 levels in both *Nf1* cKOs and *bRaf* mutants (Figure 4B, 4D WT vs. *Nf1* cKO p = 0.0004; WT and *Nf1* cHet vs. *bRaf^ca^*; *Nf1* Het vs. *Nf1* cKO p < 0.0001). Since SATB1 levels are activity-dependent, can increase SST expression when overexpressed, and directly bind to the SST promoter (Balamotis et al., 2012; Denaxa et al., 2012; Goolam and Zernicka-Goetz, 2017; Tu et al., 2019), SATB1 is a candidate for the elevated SST levels.

### SATB1 expression is elevated in *Nkx2.1-Cre*-lineage *Nf1* cKO and *bRaf^ca^* cells during development

While many of the factors probed via WB are MGE-derived and interneuron lineage at E15.5, whole forebrain was used. To validate that SATB1 protein was increased in developing CINs in the neocortex we stained E15.5 neocortices for SATB1 and found that the number of migrating tdTomato+ CINs in the neocortex expressing SATB1 was increased by 5.2 and 5.1-fold in the *Nf1* cKO and *bRaf^ca^* mutants, respectively (Figure 5A-5G p < 0.0001). These results suggest that elevated SATB1 levels in CINs derived from the MGE may be a contributor to the cell fate bias of SST+ CINs in hyperactive RAS/MAPK mutants. Moreover, it validates that MGE-derived migrating interneurons in the neocortex contribute to this phenotype.

**Figure 5:**
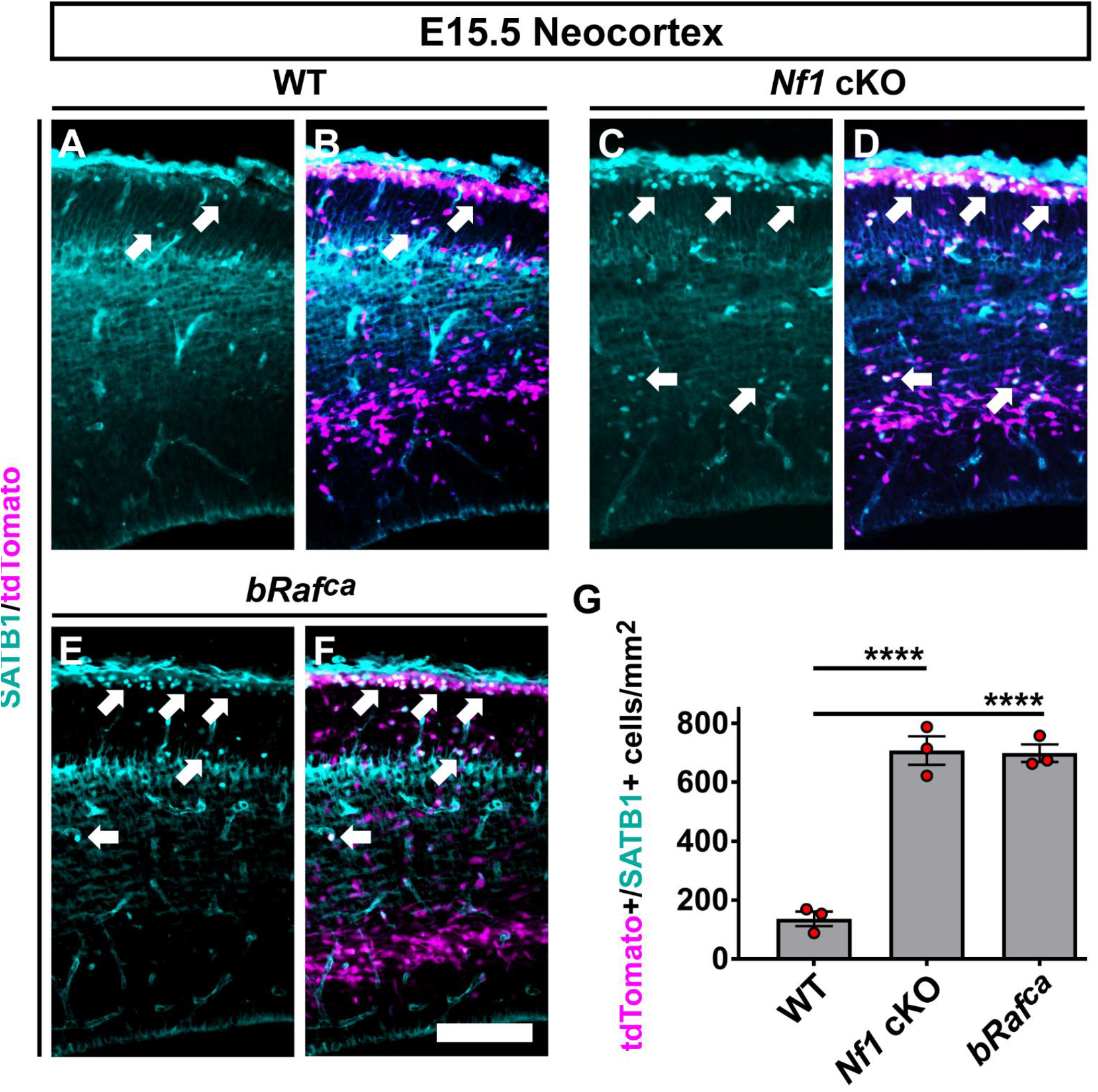
SATB1 expression is aberrantly elevated in *Nkx2.1-Cre* lineage CINs during embryonic development: (A-F) Images of SATB1 protein co-labeled with tdTomato (*Nkx2.1-Cre* lineages) in the developing neocortex. (G) Quantification of the number of SATB1+/tdTomato+ cells per area; elevated SATB1 numbers are found in both mutants. Data are expressed as the mean ± SEM, n = 3 for each group. **** p<0.0001. Scale bar in (F) = 100μm.

### ARX is decreased in both *Nf1* cKOs and *bRaf^ca^* mutants

We also assessed whether other core GABAergic CIN programs were altered in *Nf1* cKO and *bRaf^ca^* mutants. The aristaless homeobox, *Arx*, gene is one such factor but due to high expression in other cell types could not be assessed reliably via western blots. In addition to being regulated by LHX6 and DLX proteins (Colasante et al., 2008; Vogt et al., 2014; Zhao et al., 2008), it also controls CIN developmental properties (Friocourt et al., 2008; Joseph et al., 2021; Marsh et al., 2016; Ruggieri et al., 2010). We examined E15.5 brains for ARX expression and found a 31% and 44% reduction in *Nf1* cKO and *bRaf^ca^* brains, respectively (Supp. Figure 5A-G, Wt vs. *Nf1* cKO p = 0.003, WT vs. *bRaf^ca^* p = 0.0004). To determine if the loss of ARX persisted in mature CINs, we first probed for ARX in somatosensory cortices of WT and *Nf1* cKO P30 brains. ARX was decreased by 65% in *Nf1* cKO CINs (Supp. Figure 5H-K, p = 0.0003). We also assessed an equivalent age for WT and *bRaf^ca^* MGE transplanted cells. Consistent with earlier data, the proportion of transplanted CINs expressing ARX was reduced by 52% (Supp. Figure 5M-Q, p < 0.0001). Thus, ARX reduction is another shared phenotype between these two hyperactive mutants.

### Pharmacological blockade of MEK signaling normalizes SST expression in hyperactive RAS/MAPK mutants

The increase in SST+ CINs across these two distinct models suggested a link between MAPK signaling and SST expression. To test this, we employed the recently FDA-approved drug, Selumetinib, a more specific MEK inhibitor that can cross the blood brain barrier (Liang et al., 2018; McNeill et al., 2017; Van Swearingen et al., 2017). MEK activity is downstream of both *Nf1* and *bRaf* encoded proteins (Supp. Fig. 1). To test if Selumetinib could normalize SST expression, we generated MGE primary cultures from WT or *bRaf^ca^* brains and treated with either vehicle or drug every 24 hours for 8 days before assessing for SST (Figure 6A). Western blots of WT cultures treated with vehicle or 10μM or 20μM of Selumetinib were assessed for pERK to determine efficacy (Figure 6B); both drug doses were effective at reducing pERK levels, the 20μM dose was used.

**Figure 6:**
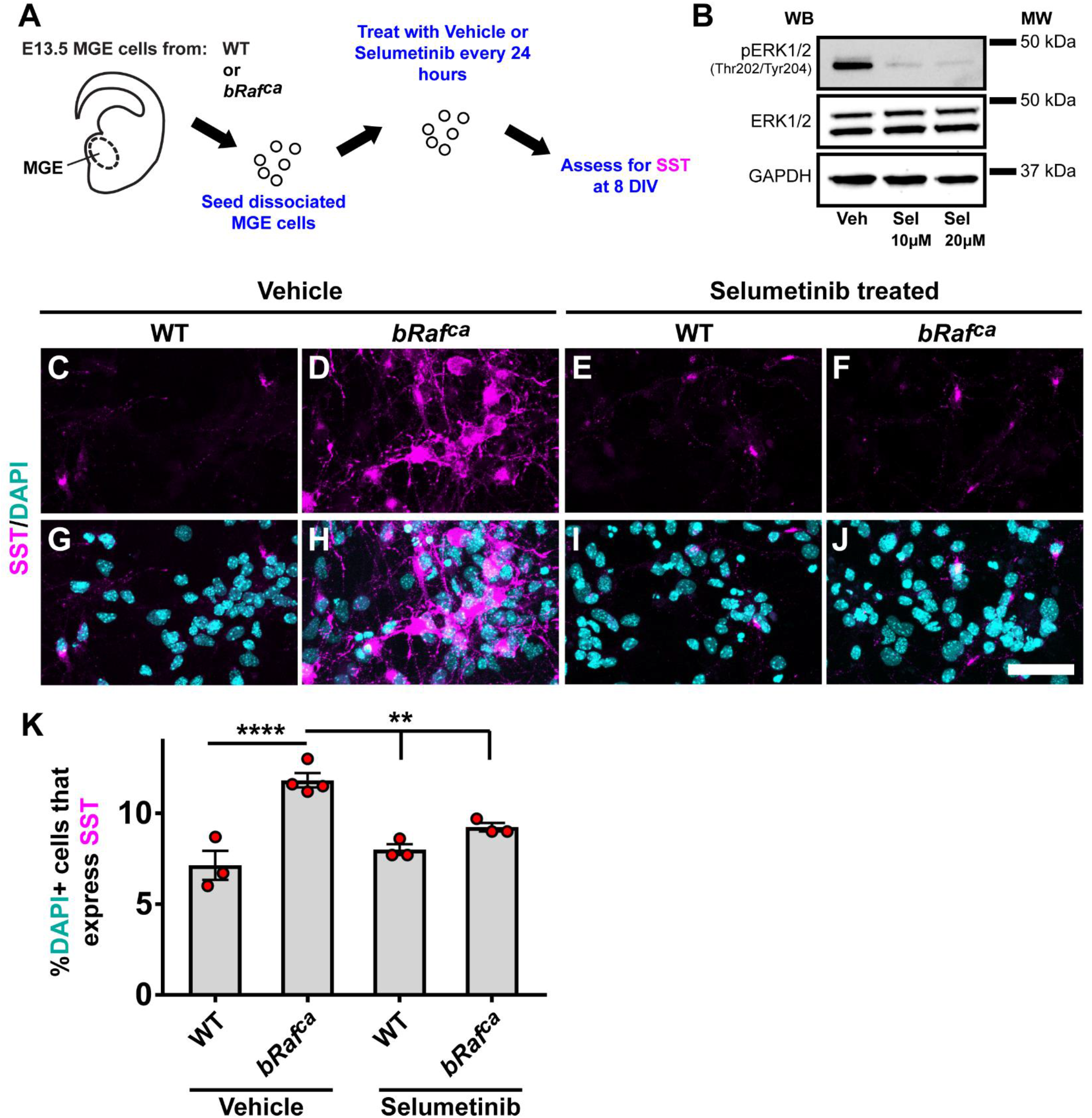
MEK inhibition prevents elevated SST and SATB1 expression in *bRaf^ca^* mutants: (A) Schema depicting the paradigm. E13.5 MGE cells were collected, dissociated and cultured in the presence of vehicle or Selumetinib for 7 days in vitro (DIV). (B) Western blots of WT cells cultured in either vehicle or drug were probed for pERK, total ERK and GAPDH at 7DIV; a 20 μM dose of drug was chosen for use. (C-R) Images of primary cultures labelled for SST, SATB1 and DAPI at 7DIV show elevated SST and SATB1 expression in the *bRaf^ca^* mutant that is prevented by drug treatment. (S, T) Quantification of the proportion of DAPI+ cells that express SST and SATB1. Data are expressed as the mean ± SEM, n = 3-4 for each group. ** p<0.01, *** p<0.001, **** p<0.0001. Scale bar in (R) = 50μm.

As expected, in vehicle treated cultures, elevated SST and SATB1 levels were observed in *bRaf^ca^* CINs (Figure 6C, D, G, H, K, L, O, P, S, SST and SATB1 p < 0.0001). 20μM Selumetinib treatment led to an attenuation of SST and SATB1 in the *bRaf^ca^* mutants but did not alter WT levels (Figure 6E, F, I, J, M, N, Q, R, T, SST, *bRaf^ca^* vehicle vs. WT drug p = 0.008, vs. *bRaf^ca^* drug, p = 0.009, SATB1, *bRaf^ca^* vehicle vs. WT vehicle p < 0.0001, vs. *bRaf^ca^* drug p = 0.0003). These data suggest that the increase in SST and SATB1 expression is dependent upon MAPK signaling.

### Activity-dependent induction of SST expression requires RAS/MAPK pathway recruitment

Finally, we sought to bring several ideas and observations together to determine how neural activity, RAS/MAPK signaling, and GABAergic core programs may work together. To this end, we used primary WT MGE cultures and allowed them to develop for 7 days with no additive or in the presence 56mM KCl as previously described (Tolon et al., 1994); KCl raises neuronal activity via depolarization and leads to a rise in SST levels. However, whether RAS/MAPK signaling is involved in this process and if this could occur in interneurons has not been tested. In addition to using KCl to raise activity in MGE cells, we also treated each group with either vehicle or Selumetinib over the course of the experiment and then assessed SST expression to determine if RAS/MAPK signaling is involved in this process. To assess if KCl addition to primary MGE neurons would result in elevated activity-dependent events, we probed for the immediate early gene FOSB using the same culture paradigm first shown in Figure 5A. FOSB expression was elevated in MGE primary cultures at the end of the paradigm (Supp. Fig. 6, p < 0.0001). Thus, our approach to elevate activity in MGE primary cultures recapitulated the rise in the activity regulated gene, FOSB, validating our premise.

MGE primary cells cultured in vehicle showed an elevated number of SST+ cells when exposed to KCl, consistent with previous literature (Tolon et al., 1994) (Figure 7A, C, E, G, I, p<0.0001). However, cells treated with Selumetinib failed to elevate SST levels when cultured in KCl (Figure 7B, D, F, H, I, p<0.0001). These data suggest that the activity-induced rise in SST levels is dependent on RAS/MAPK signaling and provide a potential mechanism for how neural activity could induce SST+ CIN properties via a powerful signaling pathway that connects extracellular cues to potential nuclear/other cellular functions (Schema, Figure 7J).

**Figure 7:**
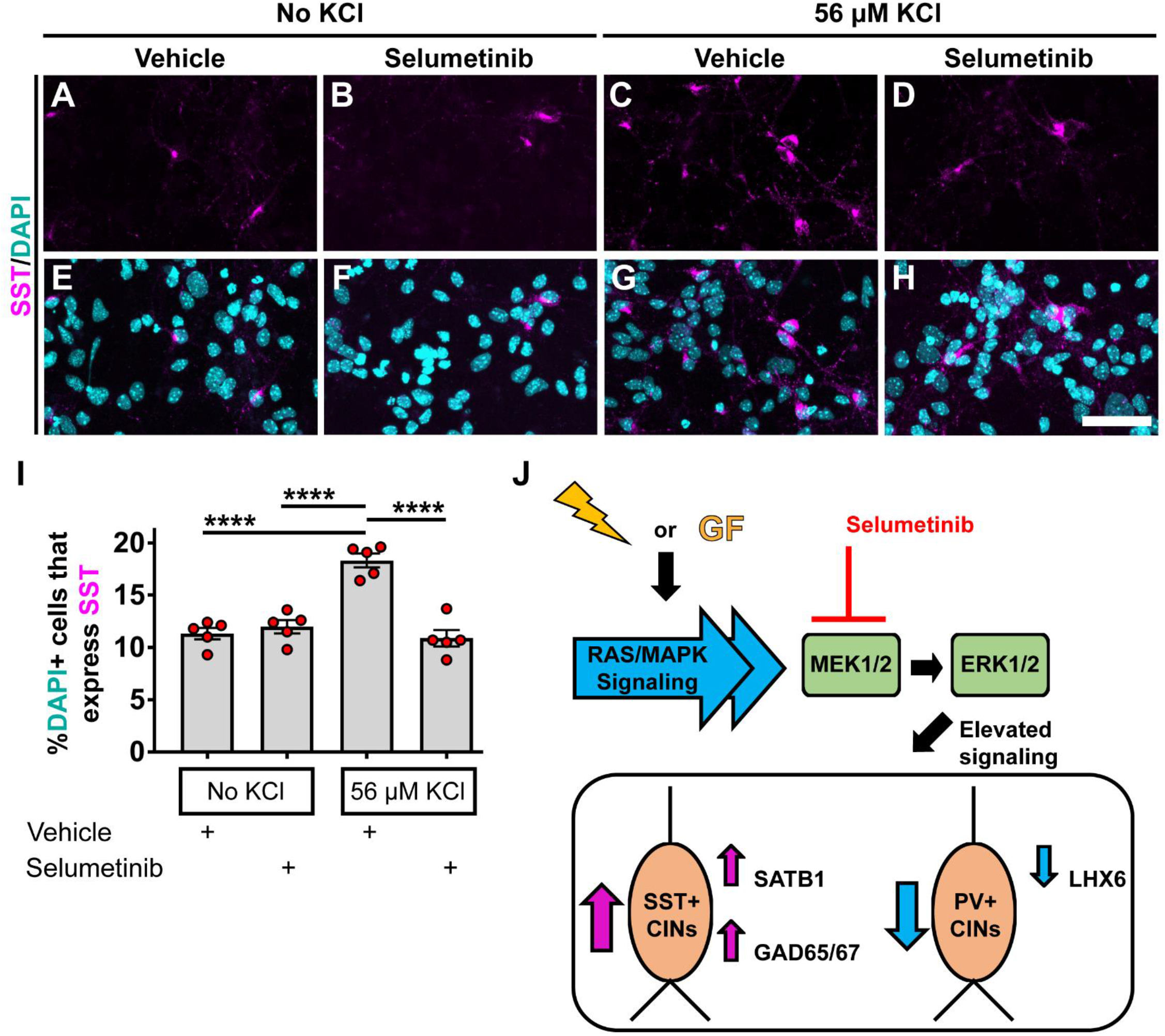
KCl-depolarized CINs utilize RAS/MAPK signaling to increase SST expression: E13.5 MGE cells were collected, dissociated and then cultured in the presence of vehicle or Selumetinib for 8DIV with each group also incubated without (A, B, E, F) or with 56M KCl (C, D, G, H). CINs were labelled for DAPI and SST after 8DIV. (I) Quantification showed that CINs incubated with 56M KCl had elevated CINs expressing SST but this phenotype was not observed in the presence of the MEK inhibitor Selumetinib. Data are expressed as the mean ± SEM, n = 5 for each group. **** p<0.0001. Scale bar in (H) = 50μm. (J) Schema summarizing the findings. Both neural activity (lightning bolt) and growth factors (GF) can recruit RAS/MAPK activity to transduce signals throughout a cell. While some RAS/MAPK proteins can signal through a variety of pathways, a shared core MAPK signaling pathway exists, which utilizes the MEK1/2 and ERK1/2 proteins. We found that RAS/MAPK hyperactive mutants also resulted in unique and common impacts on GABAergic cortical interneurons. Specifically, the bias in producing SST over PV CINs and common molecular phenotypes that include the increase in SATB1 and GAD65/67 proteins while decreasing LHX6. Our data show that these newly described phenotypes can be attenuated with the MEK inhibitor Selumetinib, suggesting these phenotypes are RAS/MAPK signaling dependent. Importantly, we show that the increase in SST expression that has been shown to be activity dependent, utilizes the RAS/MAPK signaling pathway. These data provide an important link from extracellular stimulation of CINs through canonical RAS/MAPK signaling events that impinge upon core GABAergic CIN programs.

## Discussion

We uncovered some common GABAergic CIN phenotypes caused by distinct RAS/MAPK hyperactive gene mutations. Some of these phenotypes are due to hyperactivation of the core RAS/MAPK signaling pathway. Seminal studies have pointed to the role of cardinal transcription factors in guiding interneuron cell fate and function (Liodis et al., 2007; Long et al., 2009; Sussel et al., 1999; Vogt et al., 2014; Zhao et al., 2008). Recently, neural activity and cell signaling have also emerged as important factors that guide GABAergic interneuron development and maturation (Close et al., 2012; De Marco García et al., 2011; Denaxa et al., 2012; Malik et al., 2019; McKenzie et al., 2019; Vogt et al., 2015a; Wundrach et al., 2020). Recruitment of RAS/MAPK signaling during neural activity and induction of SST expression (Tolon et al., 1994; Tyssowski et al., 2018; Wiegert and Bading, 2011; Zeytin et al., 1988) made RAS/MAPK signaling an interesting mechanism that could bridge these observation during CIN development. Our data suggest that MGE cells may bias towards SST CINs via activation of RAS/MAPK signaling, recently supported by RAS/MAPK loss of function studies (Knowles et al., 2022).

CIN development and maturation follow a well-studied timeline to produce unique cellular and molecular properties in CIN classes (Hu et al., 2017a; Lim et al., 2018; Mayer et al., 2018; Wamsley and Fishell, 2017; Wonders and Anderson, 2006). During mid gestation CINs are primarily generated in the MGE and CGE of the ventral telencephalon and after becoming post-mitotic, begin a long migration to their final cortical destinations that can be influenced by local cues and the dynamic structure of the developing brain (Fazzari et al., 2020; Wonders and Anderson, 2006). During these processes, cellular and molecular properties are starting to diverge in different CIN cell types over the course of weeks before the CINs find their synaptic partners and form the various microcircuits of the cortex. Studies have elucidated core transcription factors involved in these processes as well as the role of neural activity on these events (Batista-Brito et al., 2009; Butt et al., 2008; Close et al., 2012; Denaxa et al., 2012; Marsh et al., 2016; Pai et al., 2020; Pla et al., 2018). We found that hyperactive RAS/MAPK mutants had common changes in some of these core genes that direct development of CINs, including *Lhx6*, *Satb1* and *Arx*. LHX6 is an early determinant of MGE cell fate that is necessary for the emergence of SST and PV CINs (Liodis et al., 2007; Zhao et al., 2008) and promotes the expression of SATB1 and ARX (Denaxa et al., 2012; Zhao et al., 2008). While the loss of ARX may be through the depletion of LHX6 herein, it seems unlikely for SATB1 as expression increased in the mutants, suggesting a potential novel route of SATB1 gene or protein regulation in CINs. SATB1 is a likely candidate for the increase in SST expression in the hyperactive mutants, as previous data have shown expression of SATB1 is sufficient to induce SST expression in MGE lineages, even in *Lhx6* loss of function mutants (Denaxa et al., 2012). Overall, other core programs were not commonly altered in the hyperactive mutants, suggesting some selectivity on CIN programs regulated by RAS/MAPK activity. Future studies are needed to understand the full breadth of these changes to better understand the impact of RAS/MAPK activity on these critical cell types.

A recent idea concerning CIN development is that cortical activity influences their maturation (De Marco García et al., 2011; Karayannis et al., 2012; Wamsley and Fishell, 2017). Our data provide a potential conduit for one way this could occur. Since RAS/MAPK signaling is a prominent target of neural activity (Tyssowski et al., 2018; Wiegert and Bading, 2011), the changes we found in specific GABAergic programs could be downstream regulatory targets of activity. SATB1 expression is correlated with neural activity in developing CINs (Close et al., 2012; Denaxa et al., 2012) and we were able to increase SST levels in CINs by depolarization, as had been performed in other cells (Tolon et al., 1994). However, these observations have never been tested together. In our hands, depolarization-induced SST induction was blocked by MEK inhibition and both SST and SATB1 levels decreased in *bRaf^ca^* mutants when MEK was inhibited, demonstrating RAS/MAPK signaling is necessary. These data link extracellular effectors on CINs to developmental programs via a well characterized cellular signaling pathway. Since this pathway can be manipulated with drugs, some FDA-approved, it opens possibilities to further basic research into CIN development and maturation as well as investigating therapeutics for RASopathy cognitive symptoms.

Our data, using distinct genetic models and pharmacological manipulation, provide compelling evidence for a role of RAS/MAPK signaling in the development of CINs. However, some phenotypes may differ between unique genetic syndromes. For example, we found that loss of *Nf1* leads to elevated immature oligodendrocytes in forebrain regions (Angara et al., 2020). This was not observed in the neocortex of *bRaf* hyperactive mutants (data not shown) and could indicate other *Nf1*-regulated events (Gutmann et al., 2012) may influence ventral-derived oligodendrocytes. Despite these exceptions, the core GABAergic changes noted above do seem to be common events in the RASopathy models studied here and we predict other RASopathy models could benefit from these findings. Those RASopathy genes with ubiquitous or enriched GABAergic expression compared to excitatory cells (Ryu et al., 2019), including *Hras*, *Kras*, *Mapk1*, *Ptpn11*, *Sos1* and *Spred1*, may be of particular relevance. In turn, if common phenotypes continue to be found in additional RAS/MAPK mutants, it could also imply that shared comorbid symptoms, including ADHD, ASD and learning deficits, may be potentially treated in future studies by manipulation of GABAergic neurons.

## Methods

### Animals

All mouse lines used have been described previously. We bred *Nkx2.1-Cre* mice (Xu et al., 2008) with either *Nf1*^flox^ (Zhu et al., 2001) or *Braf^Flox-V600E^* knock-in mice (Urosevic et al., 2011), which express constitutively active (CA) *Braf^V600E^* after Cre-recombination.. Crosses included the *Ai14* (Madisen et al., 2010) Cre-dependent reporter that expresses tdTomato. *Braf* mice were initially on a C57BL/6 background and were backcrossed to CD-1 for at least three generations before experiments, to better match the genetic background of the *Nf1* mutants previously analyzed (Angara et al., 2020). In all conditions, males and females were compared but we did not find gross differences between sexes for phenotypes; biological replicates are a combination of both sexes. Experiments were approved by Michigan State University’s Campus Animal Resources and the Institutional Animal Care and Use Committee at Arizona State University.

### Electrophysiology

Mice (postnatal age: 6-7 weeks) were anesthetized with 500 μl of Tribromoethanol (Avertin) and coronal brain slices generated in in carbogen equilibrated ice-cold slicing solution containing (in mM): 110 C5H14ClNO, 7 MgCl2.6H2O, 2.5 KCl, 1.25 NaH2PO4, 25 NaHCO3, 0.5 CaCl2-2H2O, 10 Glucose, and 1.3 Na-Ascorbate. From rostral to caudal, 250 μm thick, brain slices containing S1 region of the cortex were cut using a vibratome (Leica VT1200) and incubated in solution (in mM): 125 NaCl, 25 NaHCO3, 1.25 NaH2PO4, 2.5 KCl, 1 MgCl2.6H2O, 1 CaCl2-2H2O and 10 Glucose. Incubation was performed at 34°C for 1 hour before recording (Zaman et al., 2011).

In K+-based whole-cell current clamp mode, spontaneous and evoked firing properties were recorded in tdTomato+ *Nkx2.1-Cre*-lineage CINs, in layer 1-2 of S1 region, with recording solution (32.8 ± 0.1°C) containing (in mM): 125 NaCl, 25 NaHCO3, 1.25 NaH2PO4, 2.5 KCl, 1 MgCl2.6H2O, 2.5 CaCl2-2H2O and 10 Glucose. Recording electrodes were pulled (Narishige, PC-100) from fabricated standard-wall borosilicate glass capillary tubing (G150F-4, OD;1.50 mm, ID; 0.86 mm, Warner Instruments) and had 4.3 ± 0.1 MΩ tip resistance when filled with an intracellular solution containing (in mM): 140 K-gluconate, 10 KCl, 1 MgCl2, 10 HEPES, 0.02 EGTA, 3 Mg-ATP, and 0.5 Na-GTP. The pH was adjusted to 7.35 with KOH while Osmolarity to 290-300 mosmoll-1 with sucrose. The neurons with an access resistance of 10-25 MΩ were considered for recording and the access resistance was monitored, and recordings with > 20% change were excluded from subsequent analysis. Signals were acquired at 10 KHz with a low noise data acquisition system (Digidata 1550B) and a Multiclamp700-A amplifier and were analyzed using pClamp11.1 (Molecular Devices).

### Immuno-fluorescent staining

Adult mice were transcardially perfused with Phosphate buffered saline, followed by 4% paraformaldehyde (PFA). The brains were removed and postfixed in PFA for 30 minutes. Embryonic brains were fixed in 4% PFA for 1 hour. Brains were transferred to 30% sucrose for cryoprotection after fixation, embedded in optimal cutting temperature compound and then coronally sectioned via cryostat; adult brains sectioned at 25μm and embryonic/early postnatal at 20μm. Sections were permeabilized in a wash of PBS with 0.3% Triton-X100, then blocked with the same solution containing 5% bovine serum albumin. Primary antibodies were either applied for 1 hour at room temperature or overnight at 4°C, followed by 3 washes. Secondary antibodies were applied for 1-2 hours at room temperature, followed by 3 washes and mounting with Vectashield. Primary antibodies included: mouse anti-LHX6 (Santa-Cruz Biotechnologies sc-271433), rabbit anti-PV (Swant PV27), mouse anti-SATB1 (Santa-Cruz Biotechnologies sc-376096), rat anti-SST (MilliporeSigma MAB354), rabbit anti-SST (Thermo Fisher PA5-85759, only used at P2). Secondary antibodies (used at 1:300 dilution) were either Alexa 488 or 647 conjugated and from Thermo Fischer. DAPI stained nuclei were visualized with NucBlue™ (Thermo Fisher R37606).

### Imaging

Fluorescent images were acquired using a Leica DM2000 microscope with mounted DFC3000G camera. Primary culture images were acquired using a Zeiss 800 Laser Scanning Confocal Microscope. Fluorescent images were adjusted for brightness/contrast and merged using Fiji software.

### Primary cultures

E13.5 MGE tissue was harvested and cultured as previously described (Wundrach et al., 2020). Briefly, glass coverslips were coated with poly-L-lysine, followed by laminin. MGE tissue was mechanically dissociated by trituration using a P1000 pipette tip and seeded at a density of ~200,000 cells per cm^2^. Cells were seeded in DMEM with 10% FBS serum and changed to Neurobasal containing glucose, glutamax and B27 the next day (Vogt et al., 2015b; Wundrach et al., 2020). 20 μM Selumetinib (Selleckchem S1008) was applied with new media every other day, as was vehicle (DMSO). Cells were fixed in 4% PFA on day 8 and subjected to immunofluorescent staining. Antibodies used are listed above. For KCl experiments, cells were chronically treated with or without 56mM of KCl during experiments as previously described (Tolon et al., 1994).

### Western blots

E15.5 forebrains were dissected/frozen on dry ice and then lysed in standard RIPA buffer with protease and phosphatase inhibitors and combined with Laemmli buffer (BioRad 1610737EDU) containing 2-Mercaptoethanol and incubated at 95°C for 5 minutes. Equal amounts of protein lysates were separated on 10% SDS-PAGE gels and then transferred to nitrocellulose membranes. The membranes were washed in Tris-buffered saline (TBST) and blocked for 1 hour in TBST containing 5% non-fat dry milk (blotto, sc-2324 SantaCruz biotechnology). Membranes were incubated with primary antibodies overnight at 4°C, washed 3 times with TBST, incubated with secondary antibodies for 1 hour at room temperature and then washed 3 more times with TBST. Membranes were incubated in ECL solution (BioRad Clarity substrate 1705061) for 5 minutes and chemiluminescent images obtained with a BioRad Chemidoc™ MP imaging system. Antibodies (all used at 1:4,000 dilution): rabbit anti-pCREB^Ser133^ (Cell Signaling Technologies 9198), rabbit anti-DLX2 (gift from John Rubenstein, UCSF), rabbit anti-GAD65/67 (Sigma G5163), rabbit anti-GAPDH (Cell Signaling Technology 2118), mouse anti-LHX6 (Santa-Cruz Biotechnologies sc-271433), rabbit anti-MAFB (Sigma HPA005653), rabbit anti-NKX2.1 (abCam ab76013), mouse anti-SATB1 (Santa-Cruz Biotechnologies sc-376096), rabbit anti-SOX6 (abCam ab30455), goat anti-rabbit HRP (BioRad 170-6515) and goat anti-mouse HRP (BioRad 170-6516).

## Acknowledgements

**AMS**, **KA** and **DV** were supported by the Spectrum Health-Michigan State University Alliance Corporation and the Autism Research Institute (ARI). This study was made possible by an ARI grant to **DV**. **SJK** was supported by the ARCS Foundation. **JMN** was supported by NIH grants R00NS076661 and R01NS097537. **TZ** and **MRW** were supported by NIH grants R00MH110665 and RF1MH126706. We thank Nicoletta Kessaris (University College London) and Aryn Gittis (Carnegie Mellon University) for generating and providing the *Lhx6-Cre* mouse line, respectively.

## Notes

### Competing Interest Statement

The authors have declared no competing interest.

### Summary of Updates

Reference information has been updated

