## Supplementary material for "Distinct hyperactive RAS/MAPK alleles converge on common GABAergic interneuron core programs": Supp. Figures

### Supplemental Data

#### Materials and methods

Animals: *Lhx6-Cre* [1] mice have been previously described. *Lhx6-Cre* mice were crossed to *Nf1*$^{Flox}$ and the Cre begins to express as MGE cells become postmitotic in the MGE. Other lines were documented in the main text.

Antibodies and reagents: LHX6 and SST antibodies were described in the main text. Other primary antibodies included sheep anti-ARX (R&D Systems, AF7068) and rabbit anti-FOSB (abCam, ab184938). DAPI positive nuclei were labeled using NucBlue reagent (Thermo Fisher R37606).

Selumetinib treatment: 100µM of Selumetinib dissolved in DMSO was delivered to primary MGE cultures every 48 hours; DMSO served as the vehicle control. Either vehicle or vehicle with Selumetinib was added to a stock of Neurobasal media and then distributed to individual wells of cells every 48 hours.

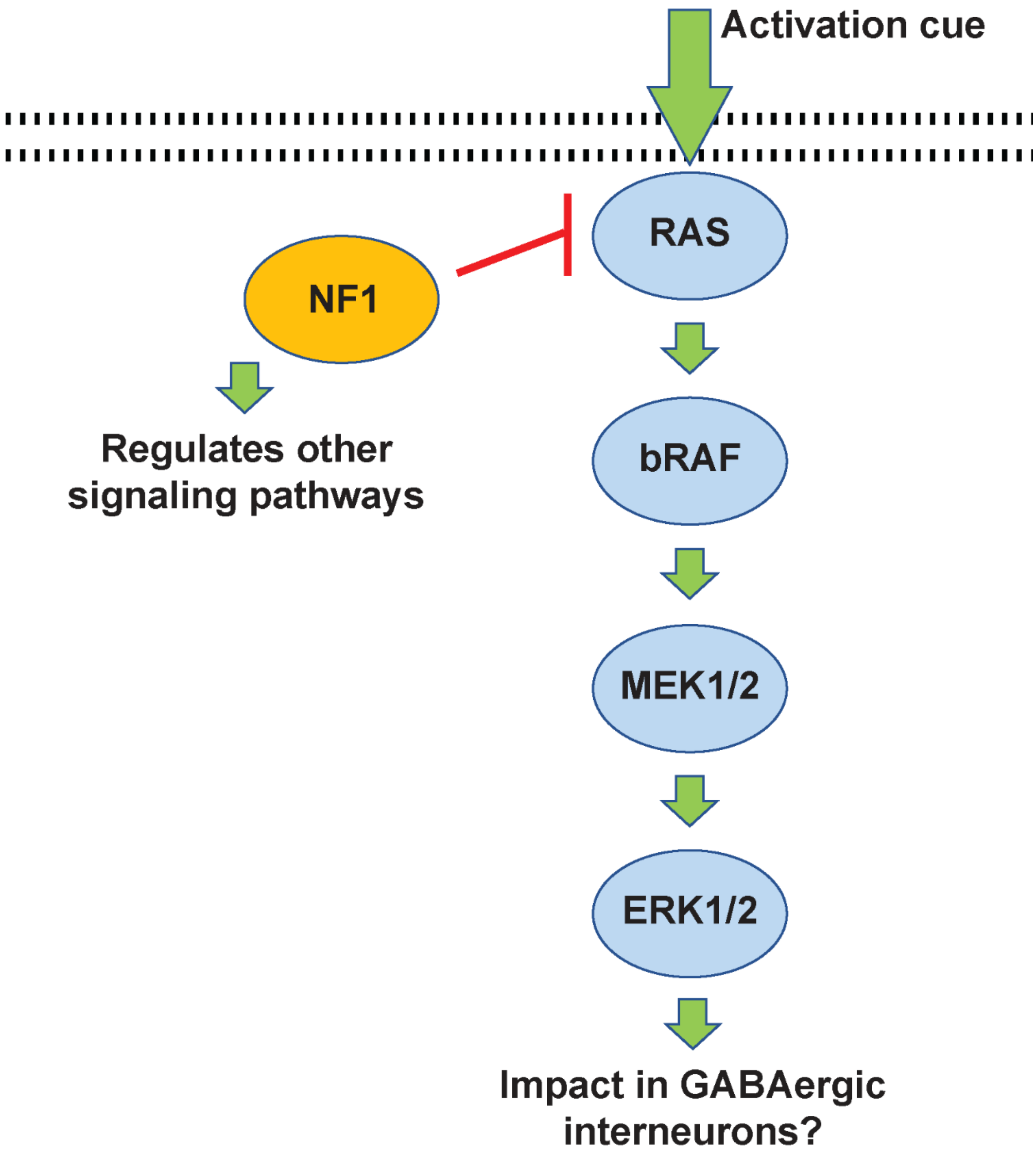


**Supplemental figure 1: RAS/MAPK signaling diagram:** The RAS/MAPK cascade is induced by extracellular signals, including growth factor recruitment and neuronal activity (activation cue). The RAS protein is the first factor activated that triggers this cascade, with RAF proteins (bRaf shown here) activated by RAS, followed by MEK1/2 and then ERK1/2 in sequence. ERK1/2 activation leads to multiple changes in the cytosol and nucleus but the impact in GABAergic cortical interneurons is not known. Note that this pathway is inhibited/shut down by NF1 activity. Since NF1 also regulates other cellular signaling events, it could influence phenotypes via multiple routes.

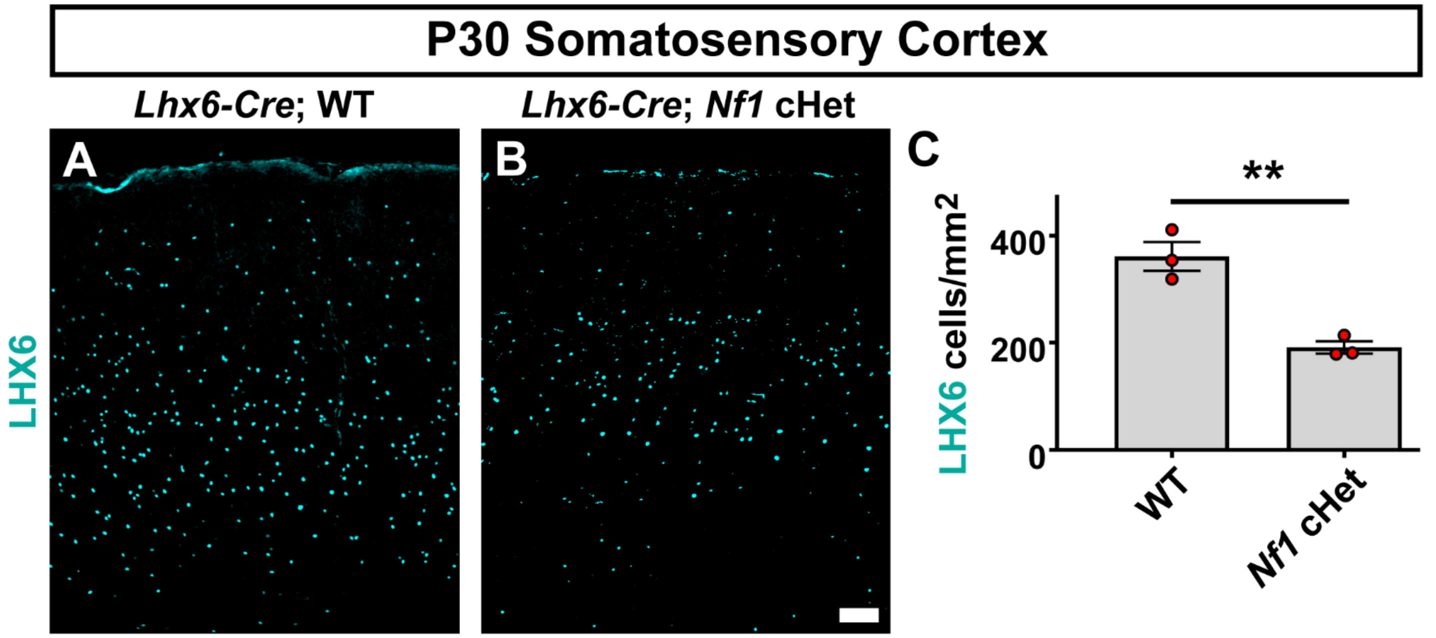


**Supplemental figure 2: Reduction of *Nf1* in *Lhx6-Cre* lineage CINs results in reduced LHX6 expression:** WT (*Lhx6-Cre* negative) and *Nf1* cHet (*Lhx6-Cre* positive; *Nf1Flox/+*) CINs were assessed at P30 for expression of LHX6 (A, B) in the somatosensory cortex. Quantification of the cell density for LHX6+ revealed a decrease, shown in (C). Data are expressed as the mean ± SEM, n = 3 for each group. ** $p<0.01$. Scale bar in (B) = 100µm.

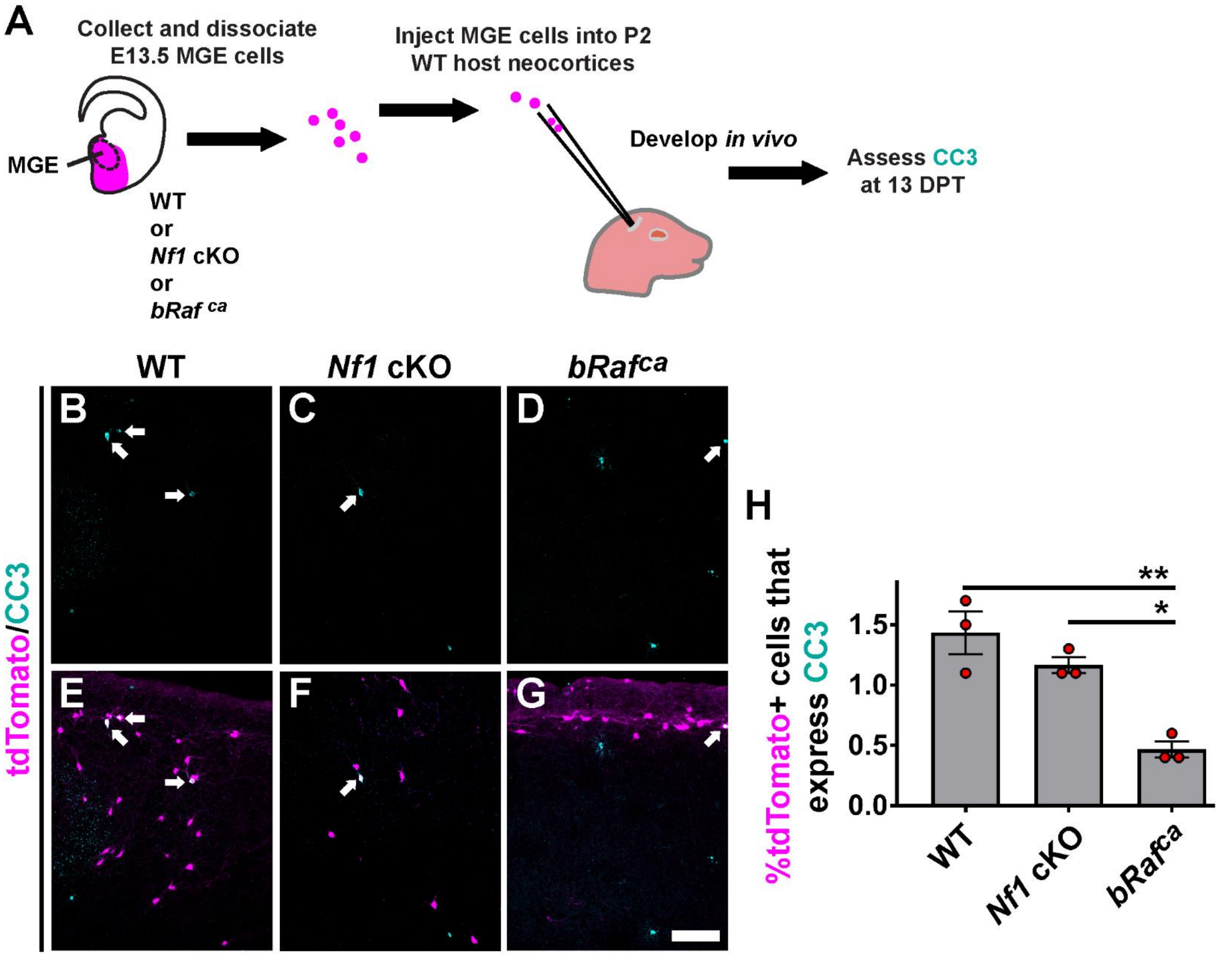


**Supplemental figure 3: Transplanted *bRaf$^{ca}$* mutant cells exhibit decreased apoptosis:** (A) Schema depicting the transplant strategy. Briefly, E13.5 MGE cells were harvested and transplanted into WT brains to develop for 13 days post-transplant (DPT), the endogenous age when apoptosis peaks. (B-G) example images of transplanted cells (tdTomato+) co-labeled for CC3. (H) Quantification of the proportion of tdTomato+ cells that co-label for CC3. Data are expressed as the mean ± SEM, n = 3 all groups. * $p < 0.05$, ** $p<0.01$. Scale bar in (G) = 100µm.

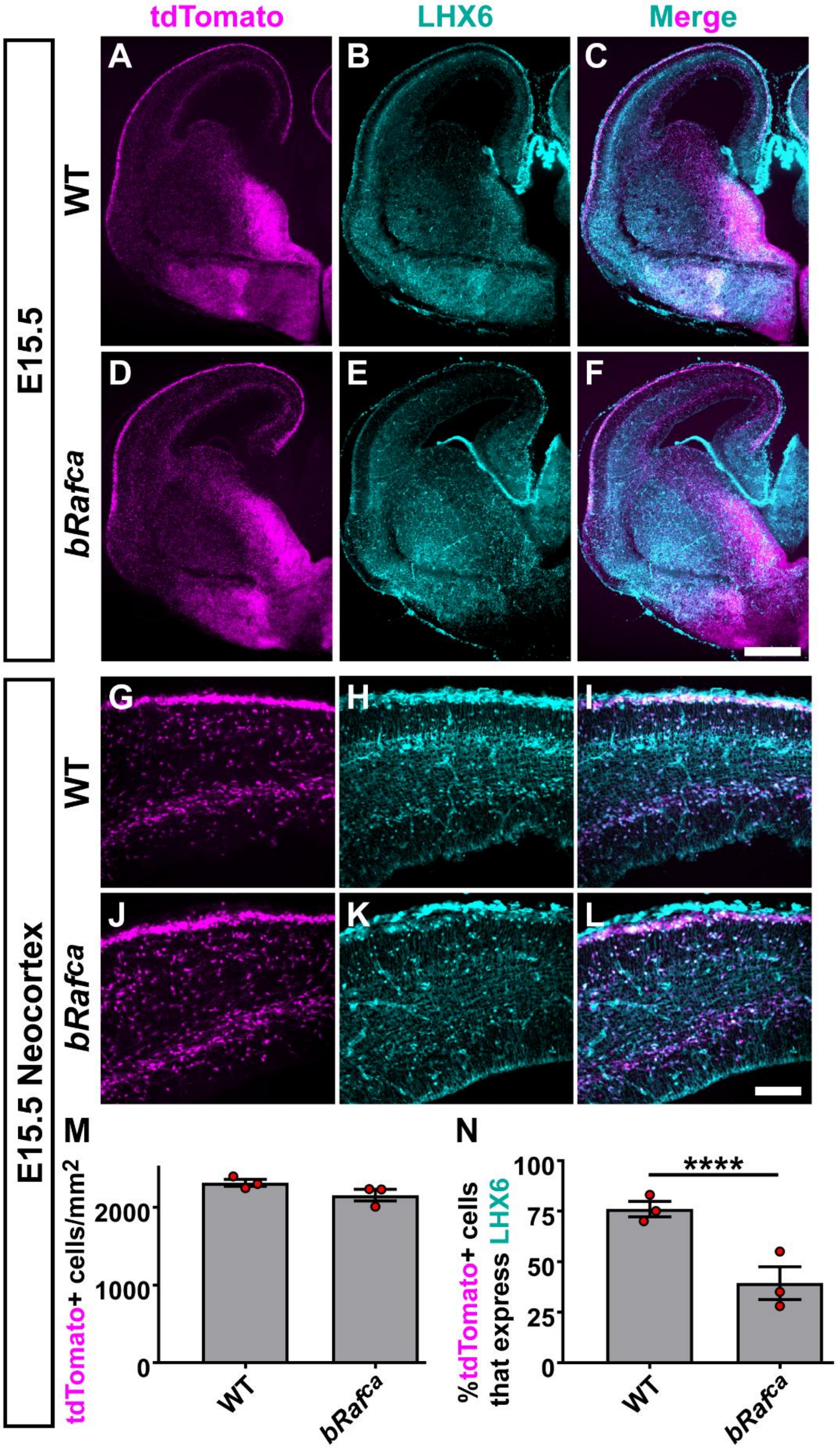


**Supplemental figure 4: Reduced LHX6 expression in *Braf^ca* neocortices at E15.5:** Coronal immunofluorescent images labeled for tdTomato and LHX6 from WT (A-C) or *Braf^ca* (D-F) embryos. Higher magnification images from the neocortex are shown for WT (G-I) and *Braf^ca* (J-L). (M) Quantification of tdTomato cell density from the neocortex reveal no change between genotypes. (N) Quantification of the proportion of tdTomato+ cells that express LHX6 from the neocortex show a decrease in the *Braf^ca*. Data are expressed as the mean ± SEM, n = 3 E15.5 brains assessed for each group. **** p<0.0001. Scale bars in (F) = 500µm and (L) = 100µm.

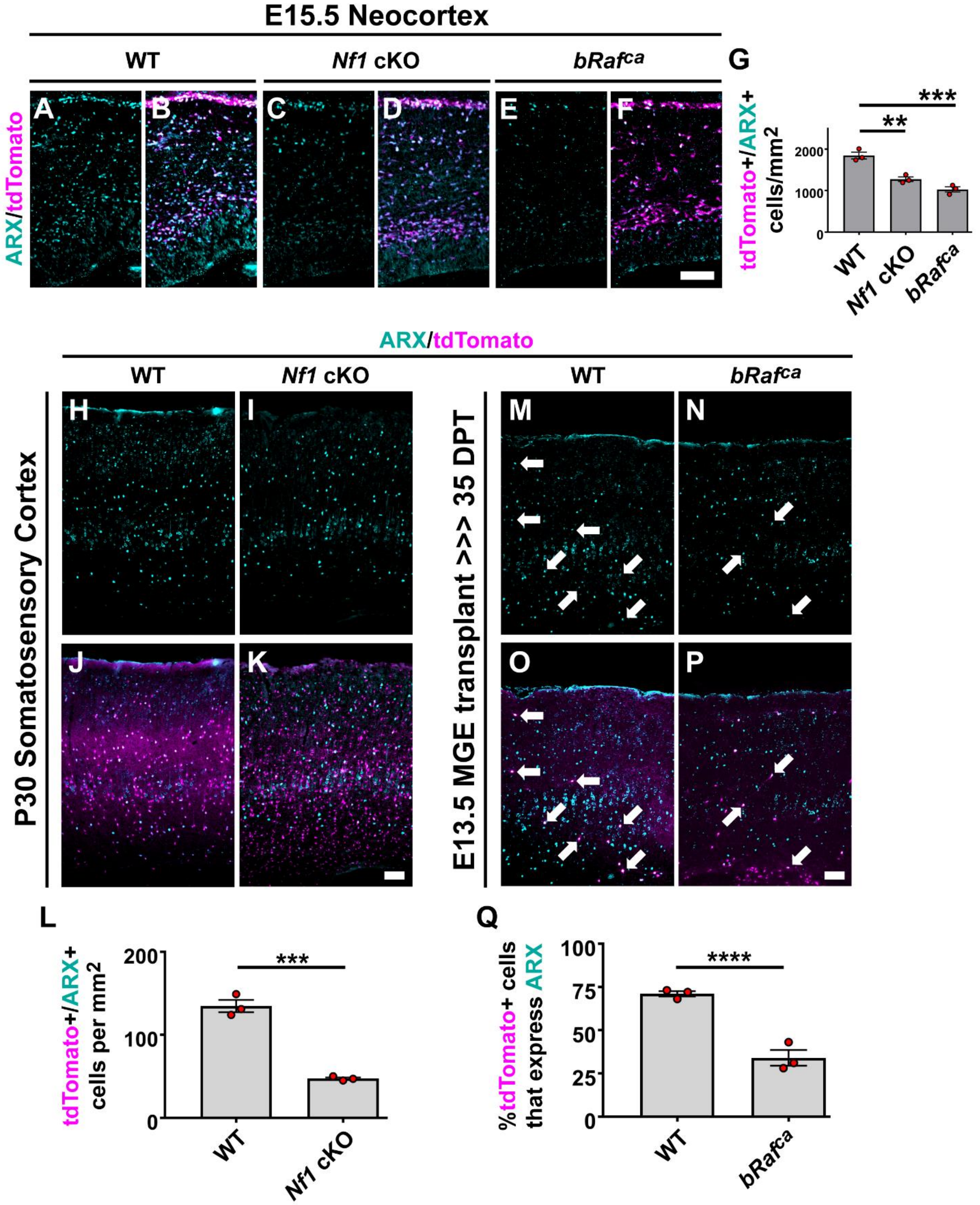


**Supplemental Figure 5: ARX is decreased in both *Nf1* cKO and *bRaf*$^{ca}$ mutants**: E15.5 neocortices were labeled for ARX protein (A-F); quantification reveals a decrease in the cell density of CINs (tdTomato+) that express ARX in both *Nf1* cKO and *bRaf*$^{ca}$ brains (G). P30 cortices were labeled for ARX expression in WT and *Nf1* cKOs at P30 (H-K), revealing a decrease in *Nf1* cKO labeled CINs (L). WT and *bRafca* E13.5 MGE transplants aged to 35 days post-transplant (DPT) were also labeled for ARX (M-P), arrows point to co-labeled cells; *bRaf*$^{ca}$ transplants also showed a reduction in ARX expressing CINs (Q). Data are expressed as the mean ± SEM, n = 3 biological replicates, all groups. ** $p<0.01$, *** $p<0.001$ and ****$p<0.0001$. Scale bars in (F, K, P) = 100μm.

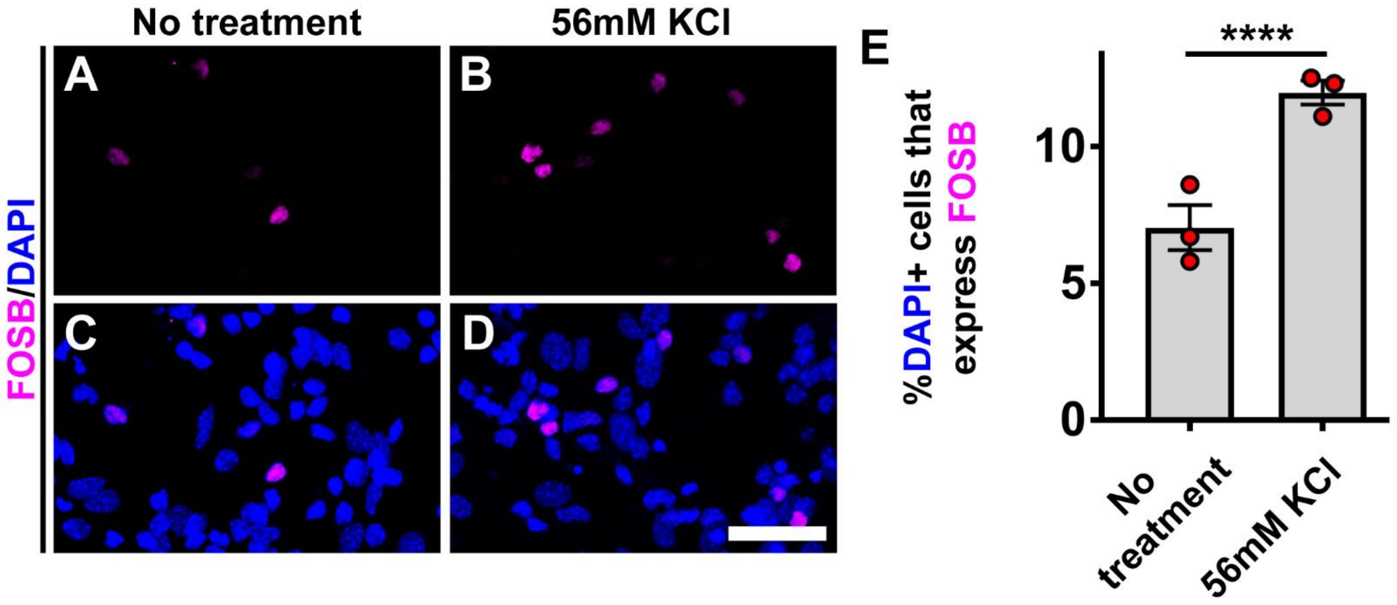


**Supplemental Figure 6: KCl treatment leads to elevated activity-dependent expression of FOSB in MGE primary neuron cultures**: (A-D) Example images of MGE primary neuronal cultures either with no treatment or in the presence of KCl for 8 days in vitro and labelled for the activity-dependent marker, FOSB. (E) Quantification of the proportion of DAPI+ cells that were labeled for FOSB after 8 days. Data are expressed as the mean ± SEM, n = 3 both groups. **** p<0.0001. Scale bar in (D) = 50μm.

| Measurements (Units) | WT | *bRaf*$^{ca}$ | p value |
|---|---|---|---|
| **Cm (pF)** | **57.4** | **64.9** | |
| SEM | ±3.6 | ±5.6 | **n.s.** |
| n | 16 | 21 | |
| **Rm (MΩ)** | **408.5** | **462.5** | |
| SEM | ±40.2 | ±5.6 | **n.s.** |
| n | 16 | 21 | |
| **Ra (MΩ)** | **16.8** | **18.3** | |
| SEM | ±1 | ±0.7 | **n.s.** |
| n | 16 | 21 | |
| **RMP (mV)** | **-64.7** | **-57.9** | |
| SEM | ±1.4 | ±1.9 | ****0.006** |
| n | 16 | 14 | |
| **Spike width (ms)** | **0.6** | **0.7** | |
| SEM | ±0.04 | ±0.06 | **n.s.** |
| n | 16 | 21 | |
| **fAHP (mV)** | **-14.5** | **-13.2** | |
| SEM | ±1.4 | ±1.5 | **n.s.** |
| n | 16 | 21 | |
| **1$^{st}$ interspike interval (ms)** | **9.9** | **18.4** | |
| SEM | ±0.6 | ±4 | **n.s. (0.08)** |
| n | 16 | 21 | |
| **Last interspike interval (ms)** | **21.9** | **49.4** | |
| SEM | ±2.3 | ±11.2 | ***0.04** |
| n | 16 | 21 | |
| **Spike frequency adaption ratio** | **2.2** | **2.8** | |
| SEM | ±0.2 | ±0.4 | **n.s.** |
| n | 16 | 21 | |

**Supplemental Table 1: Active and passive electrophysiological properties from WT and *bRaf*$^{ca}$ transplanted cells**: Chart lists measurements from transplanted WT and *bRaf*$^{ca}$ cells. Abbreviations: Cm (membrane capacitance), pF (picofarads), Rm (resting membrane capacitance), MΩ (mega ohm), Ra (active membrane capacitance), RMP (resting membrane potential, mV (millivolt), ms (millisecond), fAHP (fast after hyperpolarization).

**Uncropped western blots**

**DLX2**

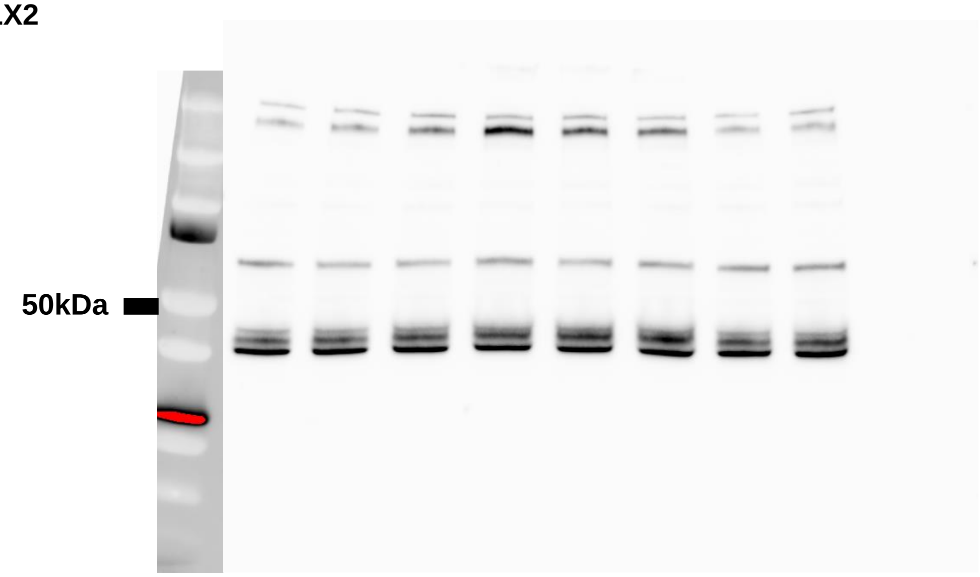


**GAD65/65**

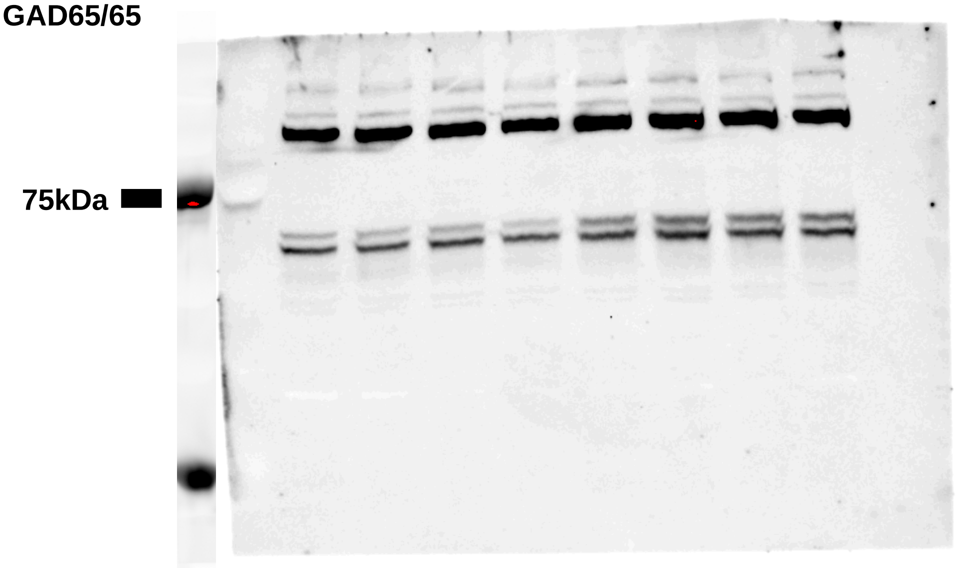


**NKX2.1**

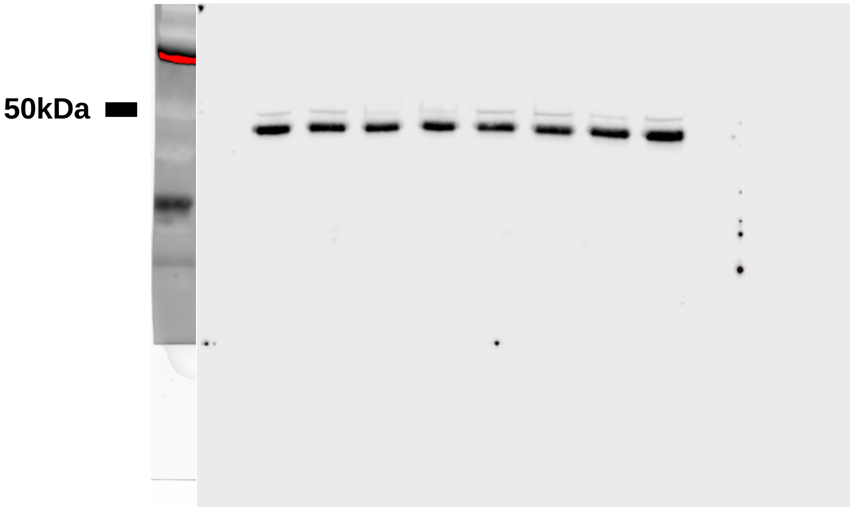

LHX6

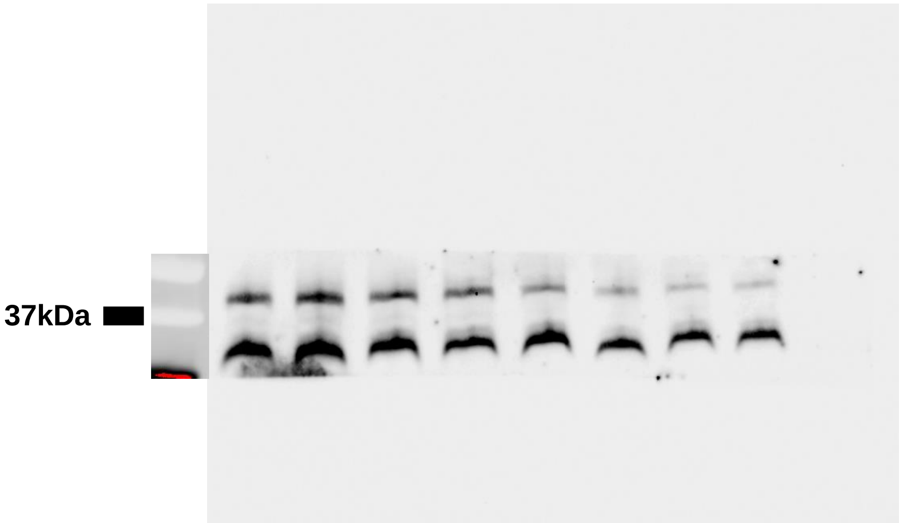


SOX6

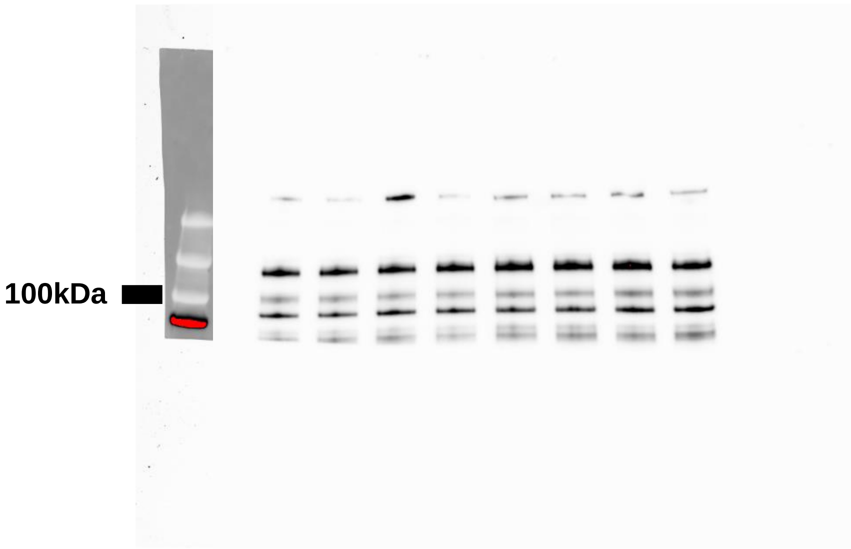


MAFB

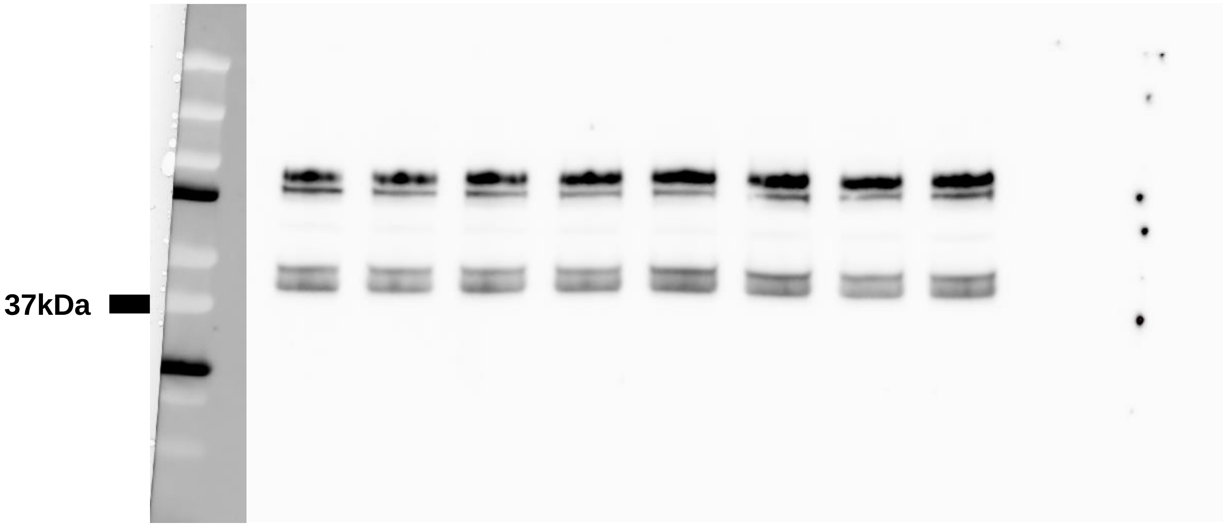

SATB1

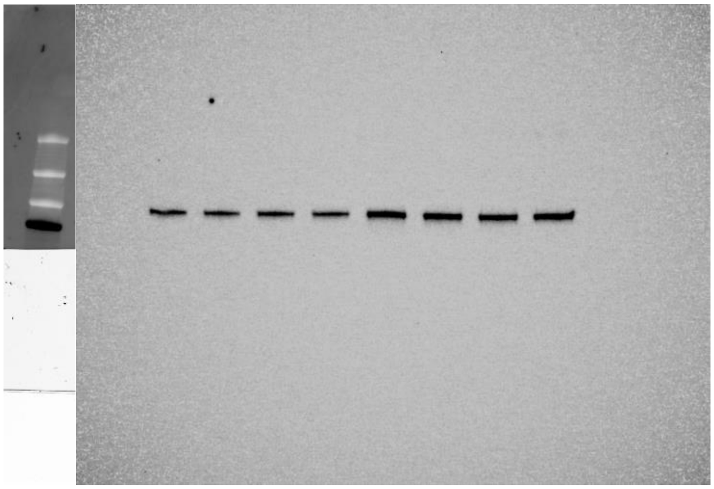


pCREB$^{SER133}$

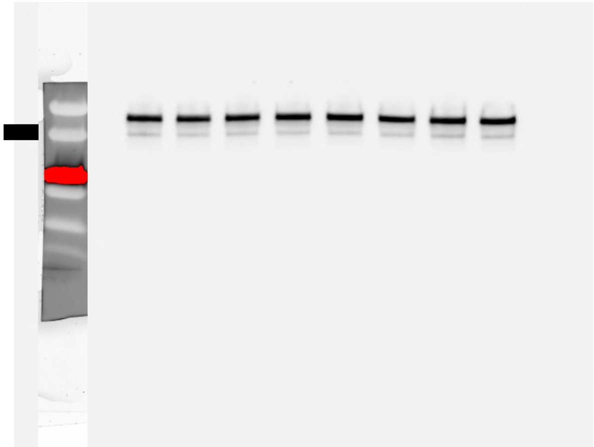


GAPDH

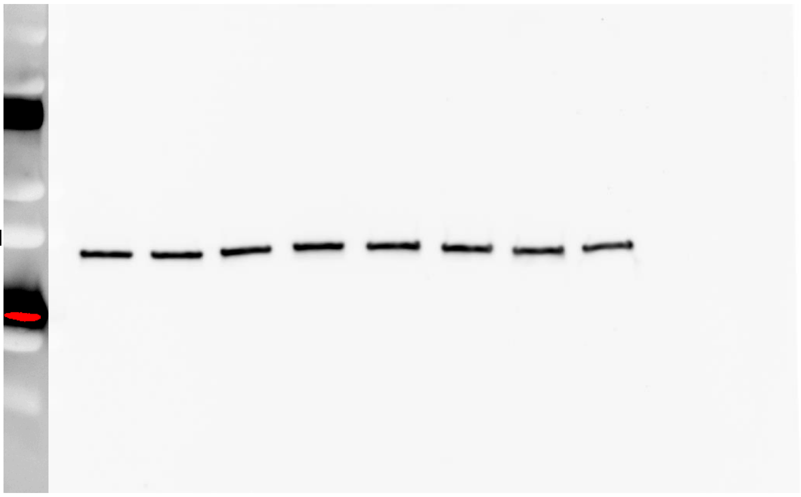
